# The Clearance of Human Cytomegalovirus Using CRISPR/Cas9 RNA Lipid Nanoparticles

**DOI:** 10.64898/2025.12.17.694960

**Authors:** Yan Ming Anson Lau, Janice Pang, Meng Qi Jiang, Yikai Sun, Karlene L.M. Knaggs, Grayson Tilstra, Ana-Maria Oproescu, Julien Couture-Senécal, Alanna M. Manning, Ranim Maaieh, Victor H. Ferreira, Atul Humar, Omar F. Khan

**Affiliations:** Institute of Biomedical Engineering, University of Toronto, Toronto, ON, Canada; Toronto General Hospital Research Institute, Toronto, Ontario, Canada; Ajmera Transplant Centre, University Health Network, Toronto, ON, Canada; Department of Laboratory Medicine and Pathobiology, University of Toronto, Toronto, ON, Canada; Department of Medicine, University of Toronto, Toronto, ON, Canada.; Department of Immunology, University of Toronto, Toronto, ON, Canada

## Abstract

Human cytomegalovirus (HCMV), a DNA virus, poses significant health risks to immunocompromised and immunosuppressed individuals and newborns. Clinical antiviral drugs like ganciclovir inhibit viral replication but have toxicities, are ineffective against drug-resistant strains, and cannot destroy HCMV DNA. CRISPR/Cas9 can cleave DNA, but has not been used therapeutically to target and degrade HCMV DNA in cells post-infection. We developed an all-in-one CRISPR/Cas9 RNA lipid nanoparticle (LNP) that clears established HCMV infections, permits rapid updating to combat resistance and is effective against multiple strains. Bioinformatic analyses identified essential, conserved viral genes as CRISPR/Cas9 targets. A delivery material screen revealed that antiviral activity was dependent on the ionizable lipid and LNP composition. Our lead formulation, βN2-40, inhibited up to 93.5% of HCMV infection with a single treatment. Furthermore, multitargeting βN2-40 LNPs demonstrated antiviral kinetics and a safety profile similar to ganciclovir, making it a compelling alternative to existing small-molecule antiviral drugs.

## Main

Human cytomegalovirus (HCMV) is a ubiquitous β-herpesvirus with an estimated global seroprevalence of 50-90%^1^. HCMV’s epidemiology is highly influenced by a given demographic’s age, socioeconomic status and country of origin^1^. This incurable, opportunistic virus is particularly detrimental to immunosuppressed and immunocompromised individuals due to their inability to control the infection^2^. These vulnerable populations include solid organ or allogeneic hemopoietic stem cell transplant recipients, HIV/AIDS patients, developing fetuses, and newborns^3–5^. For transplant recipients, HCMV infection can cause severe end-organ damage and exacerbate allograft rejection, significantly worsening transplant outcomes^2,6^. In HIV/AIDS patients, HCMV can induce retinitis and gastrointestinal complications. In severe cases, it can lead to pneumonitis and encephalitis^7–12^. In newborns, congenital HCMV infection can lead to progressive, permanent hearing and vision loss, neurological damage and other developmental complications^5^.

Ganciclovir (GCV) and its pro-drug, valganciclovir (VGCV), are the current first-line treatments for HCMV diseases^13–15^. While its antiviral efficacy is well documented, GCV and VGCV are known to cause severe myelotoxicity, most commonly characterized by leukopenia, neutropenia, and thrombocytopenia^16,17^. These hematologic complications often lead to dose-limiting measures or cessation of the therapy, which compromises the antiviral efficacy. Furthermore, administration of GCV and VGCV frequently leads to drug-resistant HCMV strains, rendering these antiviral drugs ineffective^18–20^. Other approved small-molecule antivirals, such as forscarnet and maribavir, can be used for refractory or resistant HCMV infections; however, they also suffer from toxicities and eventual drug resistance^21–23^.

Since HCMV is a DNA virus, clustered regularly interspaced short palindromic repeats (CRISPR)/CRISPR-associated protein 9 (Cas9) technologies can offer an orthogonal approach to inactivate HCMV via disruption of the viral genome. Moreover, CRISPR/Cas9’s spacer sequence can be easily modified. This adjustability can allow for quick adaptation to drug-resistant HCMV strains. The programmable nature also enables targeting of virtually the entire viral genome, expanding the number of possible antiviral targets and allows manipulation of undruggable targets. Previous reports have demonstrated some evidence of CRISPR/Cas9’s ability to attenuate HCMV replication^24–26^. However, these proof-of-concept studies used lentiviral vectors to stably transduce cells to express anti-HCMV CRISPR/Cas9 prior to infection, a strategy that is not clinically feasible. To improve the clinical potential of this approach, new systems must be developed that can treat cells post-infection without the need to constitutively express gene editors. Thus, delivering anti-HCMV CRISPR/Cas9 RNA post-infection and transiently expressing the gene editors would be an important and clinically relevant breakthrough for HCMV treatment.

Lipid nanoparticles (LNPs) have recently gained significant interest as next-generation non-viral vectors for delivering nucleic acids, particularly RNA, with implications for various diseases. Currently, there are a limited number of FDA-approved RNA LNPs, including Spikevax (mRNA-1273, mRNA LNP, Moderna), Comirnaty (BNT162b2, mRNA LNP, Pfizer/BioNTech) and Onpattro (ALN-TTR02, siRNA LNP, Alnylam). LNPs can rapidly deliver therapeutic RNA^27^, are well-tolerated^28–30^, and can be easily updated with new RNA payloads. We propose that combining the benefits of both LNP and CRISPR/Cas9 will create a new and effective solution for treating HCMV infections (Fig. 1a). To develop an effective anti-HCMV CRISPR/Cas9 LNP treatment, we used an engineering design criteria approach. A successful treatment must: (1) have broad-spectrum antiviral activity against multiple strains; (2) remain effective after infection; (3) be capable of clearing ≥90% HCMV infection with a single treatment^31–33^; and (4) provide antiviral efficacy and safety equivalent to or better than current clinical standards.

**Fig. 1:**
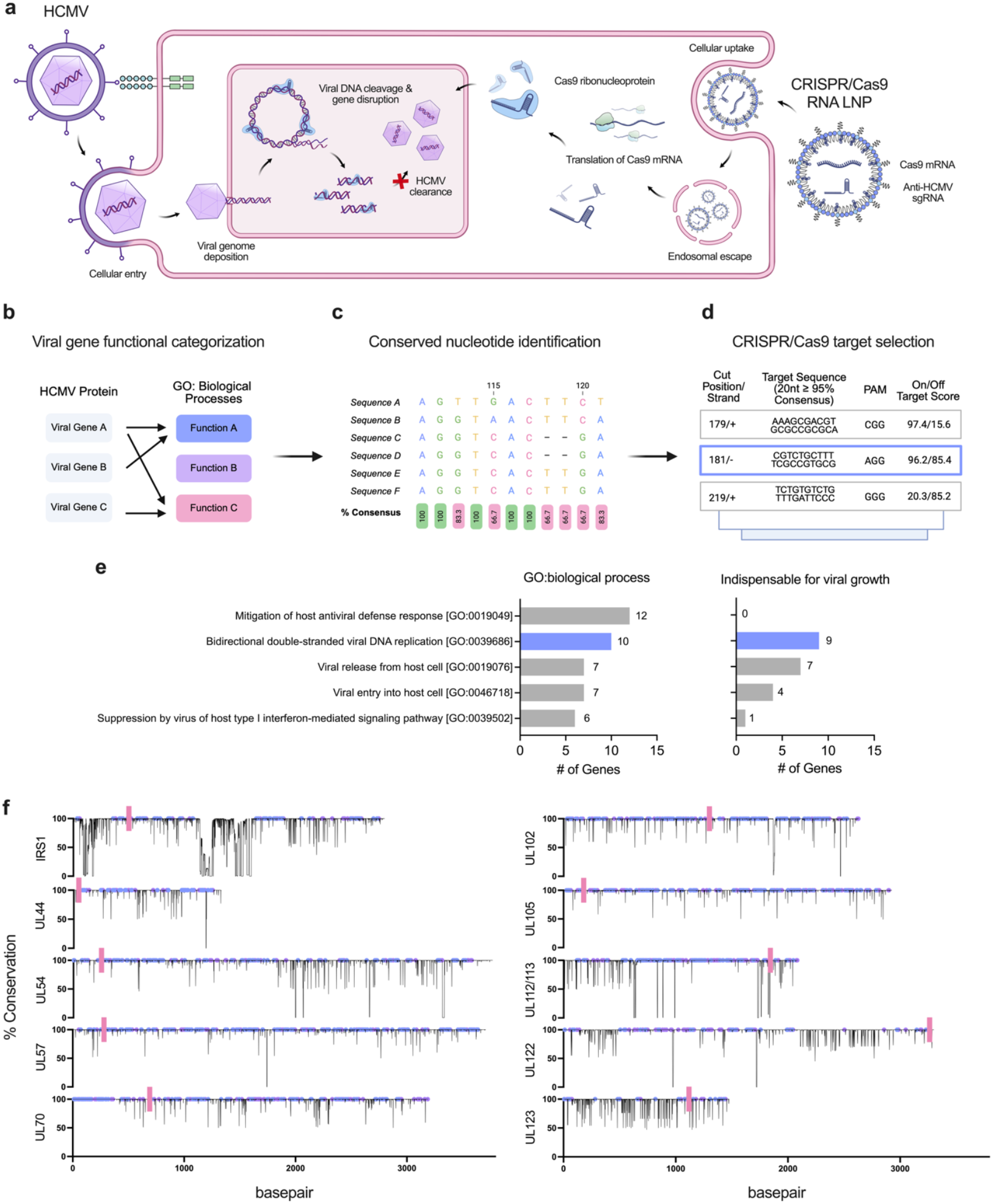
Rational selection of conserved, essential viral genes as CRISPR/Cas9 targets. **(a)** The mechanism of clearing HCMV infections using CRISPR/Cas9 RNA LNPs. **(b)** Viral gene function categorization. Viral genes were systematically groupedbased on function. **(c)** Next, all available nucleotide sequences of the categorized viral genes were aligned to identify conserved nucleotides (defined as ≥95% consensus among all input sequences). **(d)** Conserved CRISPR/Cas9 targets, defined as a (i) 20-nucleotide region with (ii) ≥95% consensus across all input sequences and (iii) an appropriate protospacer adjacent motif, that had optimal on- and off-target scores were selected. **(e)** GO: biological processes with the highest number of associated genes (top 5, left graph) and the number of essential genes in each category (right graph). Viral genes found under bidirectional double-stranded viral DNA replication classification were selected. **(f)** Nucleotide conservation plot of selected viral gene. The percent (%) conservation at a given nucleotide position was determined by DNA alignment (MAFFT) of approximately ∼346 ± 6 sequences. A conserved targetable site can be found on both the sense (blue) and anti-sense (purple) strand. The pink line depicts the location of the chosen CRISPR target.

In this report, we showed that anti-HCMV CRISPR/Cas9 RNA LNPs built to these design criteria can transiently express CRISPR/Cas9 to rapidly attenuate and provide protection against multiple HCMV strains post-infection with a single dose. We first utilized a bioinformatics approach to identify essential viral genes and conserved CRISPR target sites. Disruption of HCMV genes involved in viral DNA replication all resulted in antiviral activity, with perturbation in UL44, UL57, and UL105 resulting in the highest antiviral effects. Furthermore, a screen of different LNP formulations showed that antiviral effectiveness depended on the ionizable lipid and LNP composition, with our new formulation, βN2-40 LNP, attenuating up to 93.5% of HCMV infection with a single treatment.

## Results

### Bioinformatic selection of essential viral genes and identification of conserved targetable CRISPR sites

Deep sequencing of HCMV genomes in clinical patient samples has revealed considerable genetic variability within a single host, likely due to simultaneous multi-strain infections, de novo mutations, and recombination events between strains^34,35^. Viral gene inactivation with CRISPR/Cas9 requires complementarity between the sgRNA spacer sequence and the targeted viral DNA. However, HCMV can tolerate loss-of-function changes to its genome^36^. Thus, to identify conserved essential HCMV genes for design criterion 1, all available unique HCMV protein entries on UniProtKB (389 entries, Swiss-Prot reviewed) from different HCMV strains (AD-169, Towne, and Merlin) were categorized based on their ontological biological process (Fig. 1b). Of the 285 viral protein entries with known biological functions, 63 unique processes were identified (Supplementary Data S1). Among these processes, mitigation of host antiviral defense response (GO:0019049), bidirectional double-stranded viral DNA replication (GO:0039686), viral entry into host cells (GO:0046718), and suppression by virus of host type I interferon-mediated signaling pathway (GO:003952) were the functional subsets containing the most genes (Fig. 1e). Growth characterization compiled from previous single viral gene knockout studies revealed that all genes related to mitigation of host antiviral defense response (GO:0019049) were dispensable for viral replication (Supplementary Table S1). In contrast, genes involved in bidirectional double-stranded viral DNA replication (GO:0039686) and viral release into host cells (GO:0046718) were notably all essential for HCMV growth with one exception (Fig. 1e, Supplementary Table S1). This suggests that these biological processes are both critical for canonical viral replication and should be considered as antiviral target candidates. To reduce the risk of selecting ineffective CRISPR targets, our candidates were cross-referenced with any overlapping targets tested in previous studies where cells were stably transduced to express gene editors prior to HCMV infection ^24–26^. The viral biological process with the most validated viral genes, bidirectional double-stranded viral DNA replication [GO:0039686], was selected for further analysis.

Next, highly conserved sequences in the selected genes were identified by aligning all publicly available HCMV genomes on NCBI Virus (∼346 ± 6 sequences per gene) derived from various laboratory and clinical strains (Fig. 1c, f). CRISPR/Cas9 targetable sites were subsequently isolated. These sites were defined as (i) a 20-nucleotide region with (ii) ≥95% consensus across all input sequences and (iii) an appropriate protospacer adjacent motif (PAM, Fig. 1b). The total number of conserved, targetable sites for each selected gene and the final CRISPR spacer sequences are summarized in Table 1 and visualized in Figure 1e. A comprehensive list of all computationally determined conserved targetable CRISPR sites is available in Supplementary Data 2.

**Table 1:**
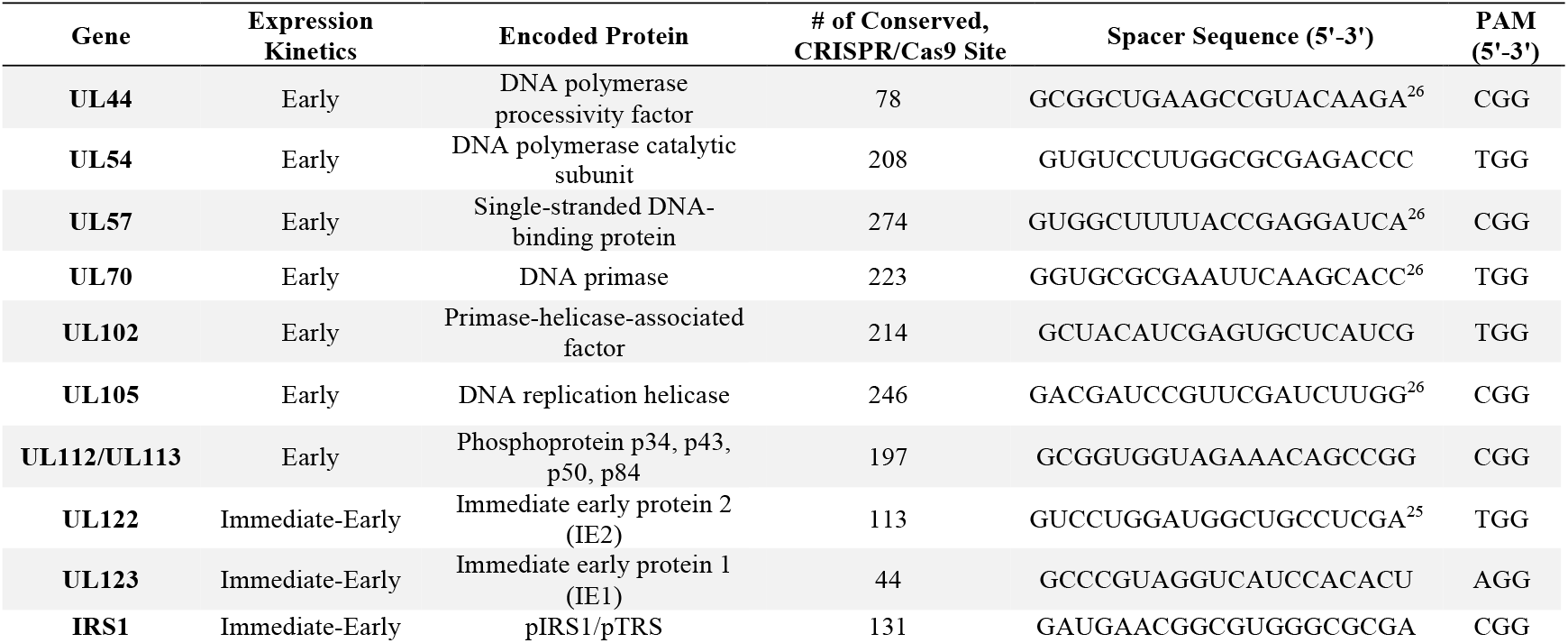
Summary of the selected viral genes, CRISPR conservation analysis, and the chosen spacer sequence against HCMV.

### CRISPR RNA LNPs attenuate HCMV replication therapeutically

In addition to inducing numerous cellular changes, including remodeled cellular architecture and arrested cell growth phases, HCMV infection can increase retention of endosomal cargo and reduce endosomal recycling events ^37–40^. Because LNP-mediated delivery requires successful endosomal escape, our next step was to confirm our ability to deliver Cas9 mRNA and sgRNA to HCMV-infected cells ^41^.

To evaluate the LNP delivery and the effectiveness of our chosen viral gene targets, SM-102 LNPs (SM-102: cholesterol: DMG-PEG2000: DSPC, mol% of 50:38.5:1.5:10) were used to deliver N1-methyl pseudouridine-modified Cas9 mRNA (mCas9) and various viral gene-targeting sgRNA (10:1 w/w) at a 20:1 lipid/RNA mass ratio. Chemically modified sgRNAs were used to improve gene-editing efficacy during infection (Supplementary Fig. S1). A pre-treatment LNP dose of 1.1 μg (RNA mass) was administered to human lung fibroblast cells (MRC-5) 6 hours prior to TB40/E-GFP infection at a multiplicity of infection (MOI) of 0.5, followed by daily LNP treatment (Fig. 2a). Flow cytometry was used to measure GFP+ levels as an indicator of infection (Supplementary Fig. S5). Six days post-infection (dpi), all CRISPR RNA LNPs targeting various viral genes significantly decreased the percentage of GFP+ cells, as compared to control LNPs containing sgRNA targeting the nonviral genomic region AAVS1 (Fig. 2b). Furthermore, the median fluorescence intensity (MFI) amongst GFP+ cells was used as a measure of viral burden per cell. All treatment groups showed a significant decrease in MFI compared with controls (Fig. 2c). Therefore, in addition to reducing the number of infected cells, CRISPR RNA LNP also decreased the viral burden per cell.

**Fig. 2:**
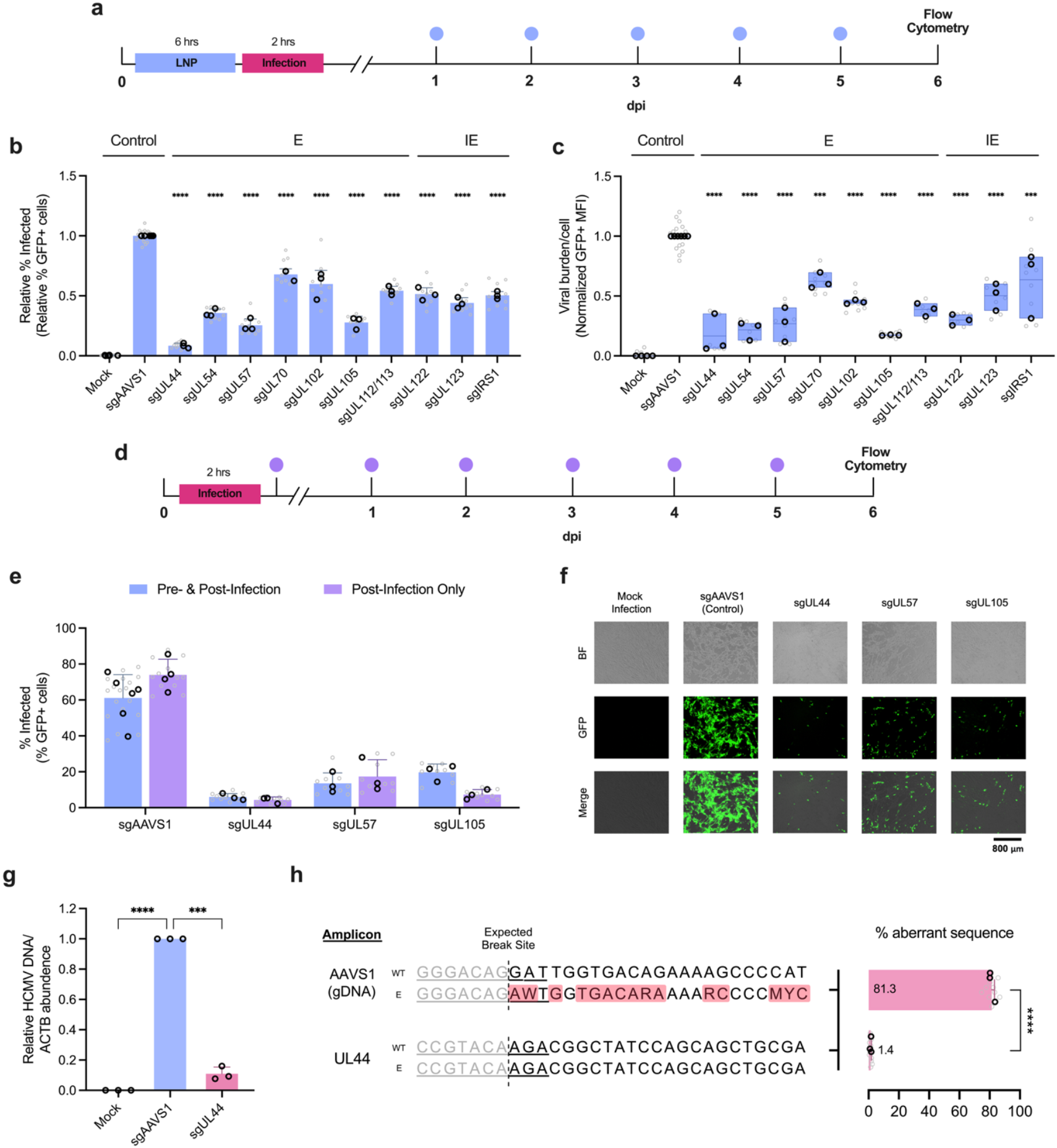
Antiviral potency of bioinformatically selected anti-HCMV CRISPR targets and their effectiveness post-infection. **(a)** Prophylactic and therapeutic treatment regimen of CRISPR RNA LNP against the TB40/E-GFP strain of HCMV.**(b)** The relative percentage of infected (GFP+) cells after prophylactic and therapeutic treatment with CRISPR RNA LNP targeting specific viral genes. Treatments were normalized to sgAAVS1 (negative control) to isolate the effects of targeted viral genes. **(c)** The normalized viral burden per cell (MFI) after prophylactic and therapeutic treatment with CRISPR RNA LNP targeting selected viral genes. Treatments were normalized to sgAAVS1. Box plots show the upper and lower bounds of infection. **(d)** Dosing regimen to treat cells post-infection. After infection with TB40/E-GFP HCMV, cells were treated with CRISPR RNA LNP. **(e)** Comparison of antiviral efficacy between pre- and post-infection dosing, and post-infection only dosing of the top-performing CRISPR targets. **(f)** Representative microscopy images of TB40/E-GFP-infected MRC-5 cells after therapeutic treatment with top-performing CRISPR targets. BF and GFP are brightfield and fluorescence images, respectively, **(g)** Abundance of HCMV DNA relative to ACTB DNA. **(h)** TIDE analysis of AAVS1 and UL44 amplicons after therapeutic treatment of CRISPR RNA LNP targeting AAVS1 and UL44, respectively. The data are shown as the mean + standard deviation, except **c**, which are shown as median + min/max. Black and gray data points denote biological and technical replicates, respectively. Statistical analyses were performed on biological replicates using one-way ANOVA for **b, c**, and **g**, two-way ANOVA for **e**, and Student’s unpaired t-test for **h**. \*\*\**P* < 0.001, \*\*\*\**P* < 0.0001.

Next, for design criterion 2, we removed the pre-treatment (prophylactic dose) and evaluated the post-infection (therapeutic) effectiveness (Fig. 2d). Previous studies showed that HCMV alters the translation landscapes in infected cells, which could hinder our ability to translate Cas9 mRNA^42^.

The top three performing CRISPR RNA LNPs from the pre- and post-infection screen were chosen (sgUL44, sgUL54, and sgUL105). Therapeutic-only treatments were as protective as the prophylactic and therapeutic combination (Fig. 2e, f). A reduction in HCMV DNA load was further confirmed by qPCR. Treatment with sgUL44 LNPs resulted in a 10-fold decrease in viral DNA compared to control LNP treatments (Fig. 2g). As an alternative method to quantify viral genome editing, we employed Tracking of Indels by DEcomposition (TIDE) analysis^43^. Surprisingly, no indels were detected at the cut site, compared to genomic DNA controls (Fig. 2h, Supplementary Fig. S2). Taken together, the findings show that LNP-delivered Cas9 mRNA and sgRNA can therapeutically inhibit HCMV replication and reduce viral load. Furthermore, inactivation of the targeted viral gene by CRISPR/Cas9 does not result in canonical indel formation, potentially indicating that cleaved viral DNA is not repaired.

### Antiviral potency is dependent on the LNP system

Next, we assessed the impact of the LNP system on antiviral activity. Our top-performing gene editing RNAs were delivered via SM-102, ALC-0315, DLin-MC3-DMA, βN2-40, and δO3-40 LNPs, with the latter two being new formulations developed by our group (Supplementary Fig. S3)^44,45^. Fibroblasts were infected with HCMV, followed by daily therapeutic treatments with different LNPs. The magnitude of antiviral activity was dependent on the LNP system, with similar trends observed across different viral targets (Fig. 3a). The most potent antiviral activity was achieved with βN2-40 LNPs targeting UL44, resulting in a 93.5% reduction in GFP+ signal compared to untreated infection (Fig. 3a).

**Fig. 3:**
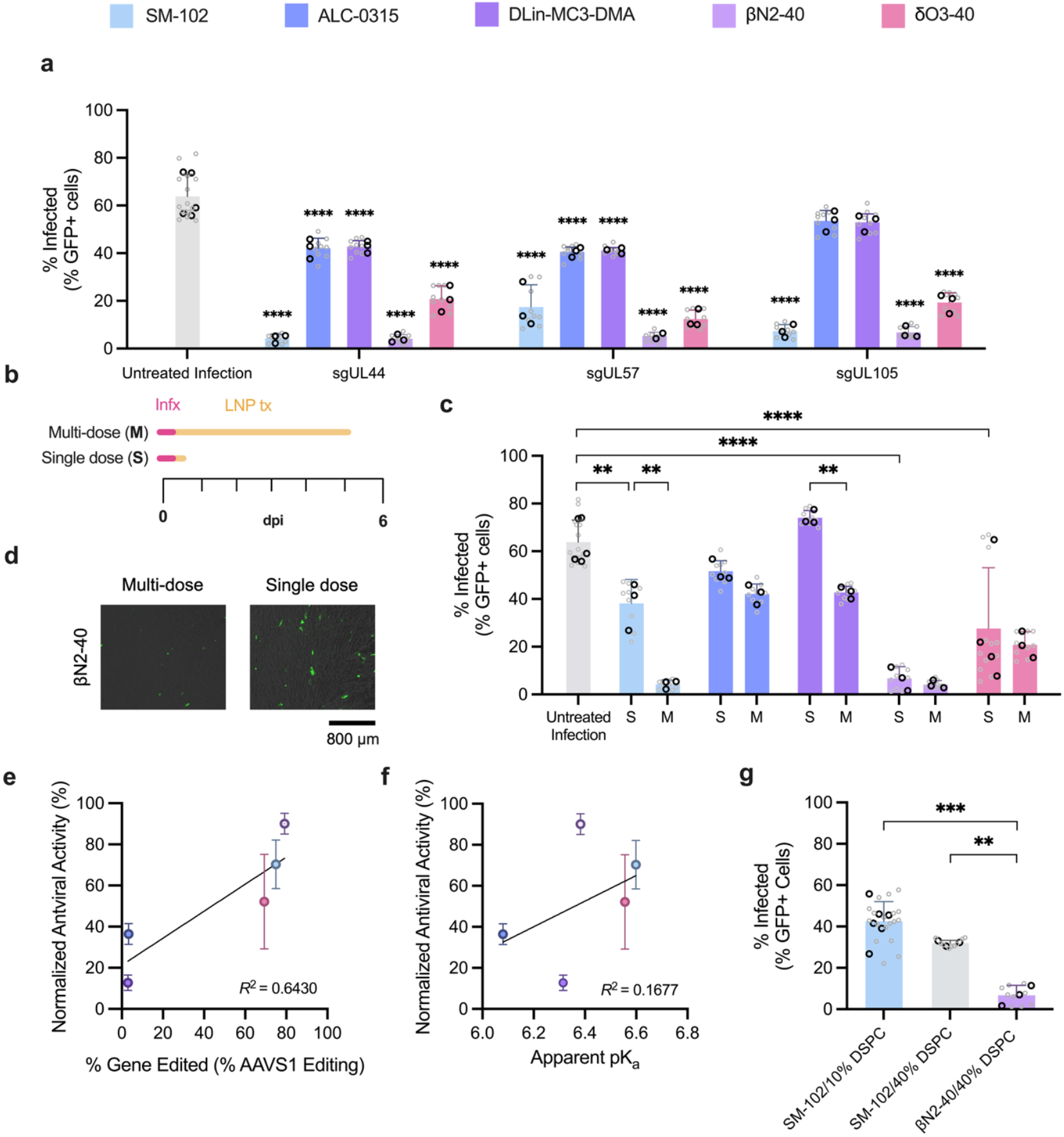
Single- and multi-dose efficacy of antiviral CRISPR LNPs is dependent on LNP composition. **(a)** Antiviral effects of top-performing CRISPR RNA delivered to pre-infected MRC-5 cells by different LNP systems, as compared to untreated infection **(b)** Schematic illustrating therapeutic multidose and single-dose CRISPR RNA LNP treatments. **(c)** Difference in antiviral effects after a therapeutic single-dose (S) or multidose (M) treatment of different LNP systems against UL44, compared to untreated infection. **(d)** Representative microscopy images of TB40/E-GFP infection after therapeutic multidose (left) and single-dose (right) treatments with βN2-40 LNPs against UL44. **(e)** Correlation between genomic DNA gene editing efficiency in healthy cells and single-dose viral DNA gene editing in HCMV-infected cells for different LNP systems. **(f)** Correlation between the apparent pKa of different LNP systems and their corresponding single-dose antiviral performance. **(g)** Antiviral effects of SM-102 LNPs, SM-102 LNPs with 40% DSPC, and βN2-40 LNPs against UL44. The data are shown as the mean + standard deviation. Black and gray data points denote biological and technical replicates, respectively. Statistical analyses were performed on biological replicates using one-way ANOVA for **a** and **g** and two-way ANOVA for **c**. \**P* ≤ 0.05, \*\**P* ≤ 0.01, \*\*\**P* ≤ 0.001, \*\*\*\**P* ≤ 0.0001.

To further resolve the differences in potency between LNPs, we evaluated the single-dose antiviral effects of each delivery system (Fig. 3b). SM-102, βN2-40, and δO3-40 LNPs targeting UL44 significantly reduced viral burden compared to natural infection after a single treatment dose, whereas ALC-0315 and DLin-MC3-DMA LNPs did not (Fig. 3c). Uniquely, a single therapeutic dose of βN2-40 LNPs was as effective as the multi-dose treatment, nearly abolishing viral growth (Fig. 3c, d). We subsequently analyzed the editing efficiency of each LNP using AAVS1 TIDE analysis. We found that the editing efficiency of non-viral genomic DNA correlated with antiviral efficacy (Fig. 3e), suggesting that effective LNP-mediated payload delivery is a critical driver of antiviral activity. To further identify the factor underlying βN2-40 LNP’s performance, we compared the apparent pK_a_ of the LNPs, a parameter that has been implicated in uptake, endosomal escape, and RNA release^46^. No correlation was found between apparent LNP pK_a_ and antiviral activity, as the top-performing βN2-40 LNP had a nearly identical pK_a_ to the lowest performer, the DLin-MC3-DLin-DMA LNP (Fig. 3f, Supplementary Fig. S4). Notably, βN2-40 LNPs also contained higher amounts of DSPC (40 mol%). To test whether the increased DSPC was responsible for βN2-40 LNPs’ potent activity, we increased the amount of DSPC in our second best-performing LNP, SM-102, to match the βN2-40 LNP’s DSPC content. A slight but non-significant increase in the antiviral activity of SM-102 was observed after increasing DSPC (Fig. 3g). This result demonstrates the critical role of the ionizable lipid for therapeutic anti-HCMV applications. The most efficient gene-editing delivery system, βN2-40, attenuated HCMV replication by >90% with a single dose.

### LNP can be multiplexed to target multiple HCMV genes simultaneously

One of the main advantages of LNPs is their ability to rapidly expand and update their therapeutic payloads. In our CRISPR antiviral context, this allows for greater target flexibility. For example, sgRNA can be quickly reprogrammed to target different sites within a single viral gene or across multiple viral genes. To assess this utility, sgUL44, sgUL54, and sgUL105 were delivered using different co-delivery strategies. These viral gene targets constitute half of the subunit types in the HCMV DNA replication complex^47^ (Fig. 4a). We expected that co-targeting different subunits in this complex would increase antiviral performance through synergy. Combinations of sgRNAs were either delivered as mixed singleplex (MS) where each sgRNA was formulated as separate LNPs and mixed at the same mass ratio as MP LNPs (Fig. 4b, center) or multiplexed (MP) LNPs, where all sgRNAs were co-formulated with equal masses of sgUL44, sgUL54 and sgUL105 (Fig. 4b, right). Singleplex (SP) LNPs containing a single sgUL44 were used as a control (Fig. 4b, left). The MP LNP formulations produced LNPs with similar physical characteristics to SP LNPs (Supplementary Table S6). Against HCMV, MS and MP LNP showed antiviral effects similar to SP LNPs across all tested delivery systems (Fig. 4c, Supplementary Fig. S6). Moreover, simultaneous disruption of UL44, UL54, and UL105 did not provide additional antiviral benefits compared to targeting UL44 alone. Critically, for design criterion 3, βN2-40 LNP remained the only LNP to achieve a >90% reduction in viral load across all delivery configurations.

**Fig. 4:**
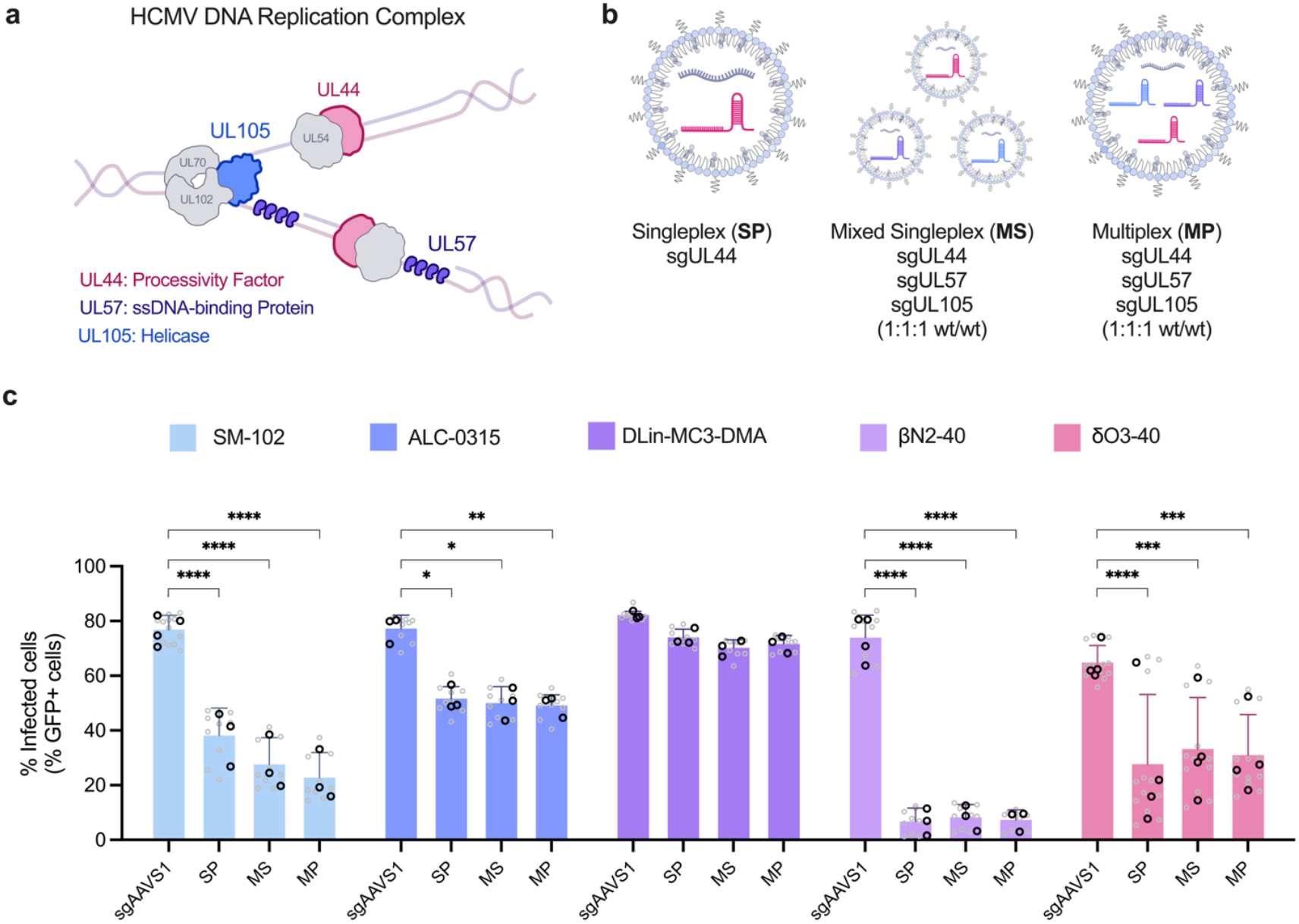
LNPs can carry multiple sgRNAs to simultaneously target different viral genes. **(a)** The HCMV DNA replication complex. The complex consists of six subunits. Highlighted subunits were co-targeted. **(b)** Schematic illustrating differences in a Singleplex (SP), Mixed Singleplex (MS), and Multiplex (MP) LNP treatment. **(c)** Differences in antiviral effects between SP, MS, and MP treatment for various LNP systems. The data are shown as the mean + standard deviation. LNPs targeting the genomic DNA target AAVS1 served as negative controls. Black and gray data points denote biological and technical replicates, respectively. Statistical analyses were performed on biological replicates using two-way ANOVA for **b**. \**P* < 0.05, \*\**P* < 0.01, \*\*\**P* < 0.001, \*\*\*\**P* < 0.0001.

### MP βN2-40 LNP inactivates HCMV with similar kinetics and efficacy as the gold standard small molecule antiviral drug ganciclovir

The MP βN2-40 LNP was selected as our lead antiviral treatment and benchmarked against CGV to address design criterion 4. Due to differences in their composition (small molecule vs RNA), we reasoned that comparing equivalent doses based on moles or mass would be uninformative. Since HCMV inhibition was the main objective, the median effective dose (ED_50_) for each therapeutic was determined as the basis of comparison. HCMV-infected fibroblasts were treated with escalating doses of GCV (0.01 – 1000 μM) and MP βN2-40 LNP (11 – 2750 ng). A sigmoidal dose curve was observed in both treatments (Fig. 5a, b). The ED_50_ for GCV in our infection model was 3.28 μM (95% CI, 2.25 to 4.88 μM; Fig. 5a), corroborating previous reports of GCV’s in vitro activity against clinical and laboratory HCMV strains^48,49^. The ED_50_ of MP βN2-40 LNP was 69.42 ng of RNA (95% CI, 57.52 to 84.42 ng, Fig. 5b). The greatest antiviral effect for MP βN2-40 was observed at 550 ng (3.95% GFP+ cells).

**Fig. 5:**
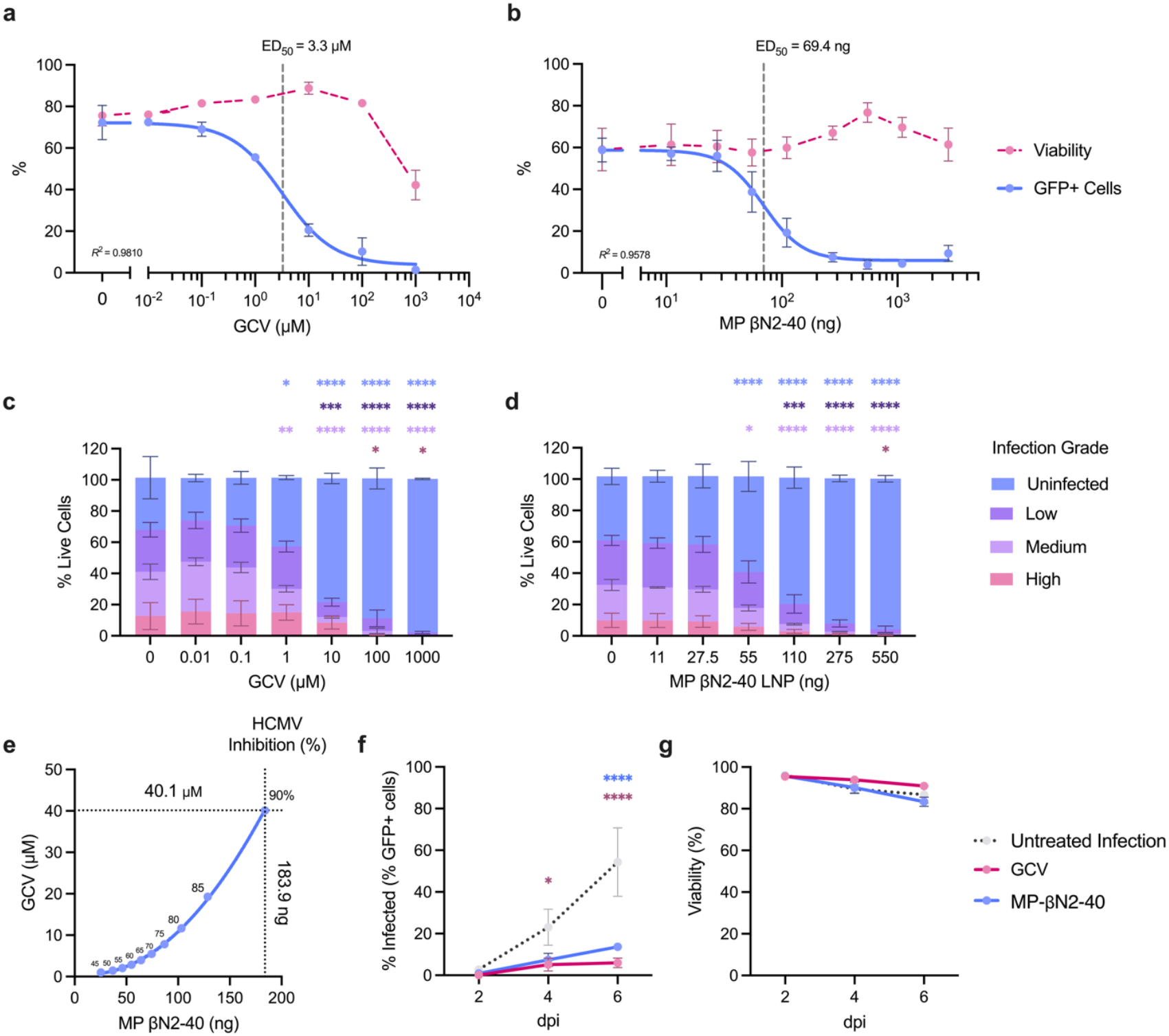
MP βN2-40 LNPs has similar antiviral characteristics as small molecule drug ganciclovir. Antiviral dose-response curve of **(a)** ganciclovir (GCV) and **(b)** MP βN2-40 LNPs and their respective median effective dose (ED_50_). The coefficients of determination of the fitted sigmoidal curves are shown. The proportion of uninfected, and low-, medium- and high-grade infection after escalating doses of **(c)** GCV or **(d**) MP βN2-40 LNPs. Infection grades were gated based on the peaks of the GFP+ histogram. Doses depicted are near the linear phase of the dose-response curve. **(e)** Predicted equivalent dose between GCV and MP βN2-40 LNP for antiviral activity. **(f**) Antiviral kinetics and **(g)** viability after GCV or MP βN2-40 LNP treatment. The data are shown as the mean + standard deviation. Statistical analyses were performed on biological replicates using two-way ANOVA for **c**, d, **f**, and **g**. For **c** and **d**, each infection grade was compared with their respective untreated control (0 μM/ng), which can be identified by the coloured legend. \**P* < 0.05. \*\**P* < 0.01, \*\*\**P* < 0.001, \*\*\*\**P* < 0.0001.

To examine the level of infection within each cell, the MFI of infected GFP+ cells was measured at different doses. We selected doses near and within the linear region of the dose-response curve. Over six days, untreated infection (0 μM/ng) showed three distinct peaks (Supplementary Fig. S8). We categorized these peaks as low-, medium-, and high-grade infections based on the histograms’ peaks and troughs (Supplementary Fig. S8). At escalating doses, GCV and MP βN2-40 LNP exhibited similar antiviral activity, with different infection grades resolving at similar dose ranges relative to their ED_50_ (Fig. 5b, c). We also observed similar inactivation kinetics across all infection grades (Supplementary Fig. S9). Interestingly, a population with high-grade infection persisted at increasing GCV doses, a phenomenon not observed with MP βN2-40 LNP treatment (Supplementary Fig. S7-9).

Using the fitted sigmodal equation, we estimated the doses required to achieve 90% HCMV inhibition to be 40.1 μM for GCV and 183.9 ng for MP βN2-40 LNP (Fig. 5e). Furthermore, when comparing the two treatments, a non-linear relationship was observed, suggesting that relatively larger doses of GCV was needed to achieve equivalent antiviral activity as MP βN2-40 LNP (Fig. 5e). HCMV-infected cells treated with either therapeutic at the predicted 90% inhibition dose demonstrated significant antiviral effects compared to untreated infections (Fig. 5f). The therapeutic onset time and viability between GCV and MP βN2-40 LNP showed no significant differences (Fig. 5f, g, Supplementary Fig. S10). Altogether, MP βN2-40 LNP and GCV have comparable inactivation kinetics, antiviral onset times and toxicity.

### MP βN2-40 LNPs targeting multiple viral genes are effective against different HCMV strains

Next, we determined if our CRISPR targets were effective against different strains of HCMV. Fibroblasts infected with TB40/E, AD-169 or Merlin strains at an MOI of 0.5, 0.5 and 0.25, respectively, were treated with MP βN2-40 LNP. Untreated infections and βN2-40 LNPs containing sgAAVS1 served as controls. Across all strains, the untreated infections showed similar viral loads to the sgAAVS1 βN2-40 LNPs controls, which demonstrated that the βN2-40 LNP itself was neither enhancing or reducing viral infections. In contrast, treatment with MP βN2-40 LNP resulted in a ∼10-1000-fold decrease in HCMV DNA across all the tested strains (Fig. 6a, b, c) (design criterion 1).

**Fig. 6:**
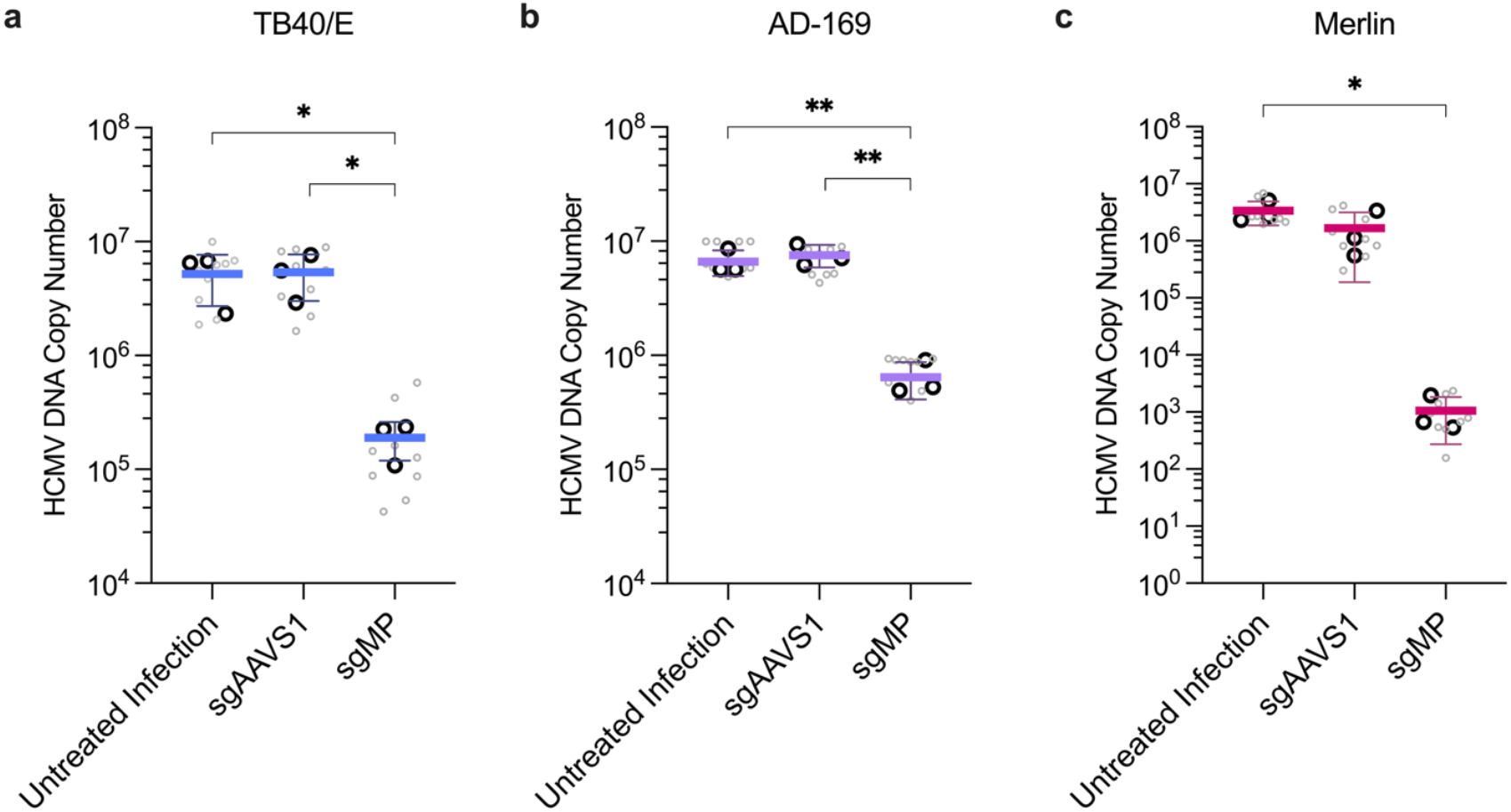
MP βN2-40 LNP provided protection against different HCMV strains. Viral DNA load of **(a)** TB40/E **(b)** AD-169, and **(c)** Merlin after MP βN2-40 LNP treatment. All cells were treated after viral infection. Black and grey data points denote biological and technical replicates, respectively. The data are shown as the mean + standard deviation of biological replicates. Statistical analyses were performed on biological replicates using one-way ANOVA for **a-c**. \**P* < 0.05, \*\**P* < 0.01.

## Discussion

We developed a broad-spectrum anti-HCMV therapeutic platform comprised of CRISPR RNA and the βN2-40 LNP. This platform allows transient expression of CRISPR/Cas9 in HCMV-infected cells and permits rapid addition of CRISPR targets. Our lead MP βN2-40 LNP demonstrated antiviral activity and tolerability comparable to the current clinical standard, GCV, suggesting its clinical potential as a new antiviral modality. Furthermore, this work is the first example of an all-RNA CRISPR/Cas9 approach to therapeutically treat HCMV infections.

This outcome was achieved through an engineering design criteria-based approach. Guided by our first criterion of creating an antiviral that is effective against all strains of HCMV, we utilized a bioinformatics approach to identify critical viral genes and conserved CRISPR-targetable sequences that were applicable across hundreds of HCMV strains. This strategy provided an efficient, rational selection process for identifying anti-HCMV CRISPR targets that are effective, broad-spectrum, and less likely to be affected by viral mutations^50^. All bioinformatically selected CRISPR targets demonstrated significant antiviral effects and across different HCMV strains, showing the utility of this method.

Of all tested targets, UL44 (DNA processivity factor), UL54 (DNA polymerase catalytic subunit), UL57 (DNA-binding protein), and UL105 (helicase) disruption resulted in the highest level of HCMV inhibition. These viral genes, along with UL70 (primase) and UL102 (primase-associated factor), form the lytic origin of replication (oriLyt)-dependent DNA replication complex^47^ (Fig. 4a). Disruption of these subunits individually resulted in varying levels of antiviral activity, suggesting differences in loss-of-function tolerances. GCV’s antiviral effect is also exerted through this complex by competitively inhibiting UL54 and inducing premature chain termination during viral DNA synthesis^51^. Disrupting UL44, UL57, and UL105, which are currently undruggable, yielded greater antiviral activity than UL54. Interestingly, targeting UL44, UL57, and UL105 simultaneously produced identical antiviral effects as targeting UL44 alone. Our data shows that, for a given viral genome, cleaving multiple sites appears to have the same effect as efficiently cleaving a single site. Thus, the multiplexing capability is best utilized by potently, targeting one effective viral gene per strain instead of multiple genes in a single strain.

Unexpectedly, ablation of the IE genes UL122 and UL123 did not result in the greatest antiviral effects. The first proteins expressed during lytic infection are produced from UL122/UL123, IE86 and IE72, which are the central initiators and regulators of HCMV’s coordinated viral gene expression cascade^52,53^. Due to their crucial role, these IE proteins have been implicated as strong antiviral candidates^25,24,54,55^. However, our data suggests that targeting IE1 and IE2 is less effective than targeting subunits within the oriLyt-DNA replication complexes at equivalent doses (Supplementary Fig. S11). This implies that the IE proteins may be suboptimal antiviral targets, perhaps due to a low IE1/2 requirement for initiating the viral gene cascade.

The need to create an antiviral modality that is relevant to clinical use also led to our second design criterion of post-infection treatment efficacy. This design specification challenged us to rely on a delivery system that can efficiently deliver the CRISPR RNA payload while inducing minimal toxicity, especially during a vulnerable, infected state. When anti-HCMV CRISPR/Cas9 RNA was administered post-infection by Lipofectamine, it failed to stop HCMV infection and caused significant cell death after a single treatment (Supplementary Fig. S10). In contrast, βN2-40 LNPs delivered anti-HCMV RNA payloads with minimal toxicity and potently attenuated HCMV. These results support the conclusion that the delivery system is critical for turning CRISPR RNA into antiviral therapeutics.

Our third design criterion aimed to maximize antiviral effect while being dose sparing. We set our minimum antiviral efficacy threshold at 90%, as this value corresponds to GCV and VGCV’s clinical efficacy and positive patient outcomes^31–33^. Our delivery screen showed that the best antiviral LNPs for post-infection cells were the ones that had the greatest genomic DNA editing efficiency in healthy cells (Fig. 3e). Amongst the tested LNP formulations, only βN2-40 LNP met and exceeded our single-dose 90% inhibition threshold, achieving 93.5% inhibition (Fig. 3c and 4b). We hypothesized that βN2-40’s superior performance was due to its pK_a_ and increased phospholipid content. However, this hypothesis was rejected due to the lack of an observable relationship between LNP pK_a_ and antiviral activity (Fig. 3F). Additionally, increasing the phospholipid content of the second-best performing LNP, SM-102, to match that of the top performing βN2-40 LNP failed to produce significant improvements (Fig. 3g). Thus, the ionizable lipid plays a critical role in facilitating a strong antiviral effect. The major molecular differences between βN2 and SM-102, which may have contributed to its improved performance, are three tertiary amines, symmetric tails, and amide resonance structures (Supplementary Fig. S3).

Our fourth design criterion was to achieve equivalent performance to current clinical gold standards. When comparing the performance of MP βN2-40 LNP and GCV, we observed similar antiviral activity, kinetics, and therapeutic onset times (Fig. 5). This result was unexpected because the two therapeutics have very different initiation and inactivation mechanisms. CRISPR RNA requires translation and complexation to generate functional antiviral units that disrupt viral replication by DNA cleavage. Past expression kinetic studies of RNA LNPs showed that protein translation can begin within one hour^56,57^. In contrast, GCV requires viral kinase UL97 and cellular kinases for tri-phosphorylation so it can bind to UL54 and inhibit viral replication^58^. GCV triphosphate, the active antiviral form, can take up to 10 hours to become detectable in infected cells^59^. In addition to performance similarities, we also observed comparable safety profiles for MP βN2-40 LNP and GCV.

While analyzing the effects of CRISPR’s antiviral activity, we discovered that CRISPR cleavage did not result in indel formation within the HCMV genome despite a measurable reduction in viral DNA and mature virion load. This observation was unexpected as genomic DNA editing often leads to the generation of indels at the cleavage site, facilitated by non-homologous end joining (NHEJ) repair mechanism^60^. Our observation suggests that viral DNA does not recruit or interact with cellular host DNA damage response (DDR) elements required to initiate NHEJ^61^. This aligns with previous reports that Ku70/80 and DNA-dependent protein kinase (DNA-PK), proteins required for NHEJ, are excluded from HCMV viral replication centers^62–64^.

Therapeutic interventions are essential to rapidly address the health risks caused by infectious diseases. The CRISPR/Cas9 system evolved in prokaryotes as a defence mechanism against viral infection^65^. We leveraged evolution to create a therapeutic that is orthogonal to current small molecule drug approaches. With appropriate RNA design and optimized delivery technology, we have shown that these gene editors can be therapeutically deployed against multiple human virus strains post-infection and can be rapidly updated to combat new viral targets. This approach holds great promise for the clinical treatment of incurable HCMV infection.

## Methods

### Viral gene function analysis and conserved CRISPR target identification

389 HCMV protein entries on UniProtKB (Swiss-Prot reviewed) from three different HCMV strains (AD-169, Towne, and Merlin) were categorized based on their ontological biological process. Viral genes within the most abundant groups were cross-referenced within the literature to determine their essentiality. Subsequently, all available nucleotide sequences of the selected viral genes were obtained from NCBI Virus (Virus/Taxonomy: Cytomegalovirus humanbeta5, taxid:3050295, nucleotide completeness: complete, host: Homo sapiens [human], Taxid: 9606) and aligned using Geneious Prime (MAFFT) to determine % conservation at a given nucleotide position. From here, conserved CRISPR/Cas9 targetable sites were identified, defined by a continuous 20-nt region that possessed ≥95% consensus across all input sequences with an appropriate protospacer-adjacent motif.

### RNA preparation

Codon optimized Cas9 sequence with T7 sequence, optimized 3’ and 5’ untranslated region, and a 101 polyA tail were cloned into a pUC57-Kan-mini for mRNA production. Modified Cas9 mRNA was produced via in vitro transcription (4 hours) in the presence of N1-methylpseudourdine nucleoside and co-transcriptionally capped with CleanCap® Reagent AG (New England Biolabs, E2080). In vitro transcribed Cas9 mRNA was purified using RNA cleanup spin columns (New England Biolabs, T2050), resuspended in nuclease-free water, and stored in −80 °C until use. Modified sgRNAs (Sythego), prepared with a 2’ O-methyl analog on the first and last three bases, and 3’ phosphorothioate antinucleotide linkages between the first three and last two bases, were used.

### CRISPR/Cas9 LNP synthesis

Ionizable lipid (βN2^44^, δO3^45^, SM-102 (Cayman Chemical, 33474), ALC-0315 (Cayman Chemical, 33474) or DLin-MC3-DMA (Cayman Chemical, 34364), cholesterol (VWR, 0433), DSPC (TCI, D3926), and DMG-PEG 2000 (Avanti Research, 8801510) dissolved in ethanol were rapidly mixed with modified Cas9 mRNA and sgRNA (10:1, wt/wt) in 25 mM, pH 4.5 sodium acetate (Thermo Scientific, J63669) in a PDMS-casted microfluidics channel. To mix, a flow rate ratio of 2.5:1 (aqueous: organic) with a total flow rate of 4.2 mL/min was used. The mixed product was directly flowed into a 10K MWCO dialysis cassette (Thermo Scientific, 66005) and underwent buffer exchange into 20 mM Tris-HCl (Invitrogen, 15567027) overnight. Subsequently, LNPs were sterile filtered (0.22 μm) (Cytiva, 99161302), adjusted to 8% sucrose, and stored in −80°C. Particle size, PDI, and zeta potential were measured by dynamic light scattering using a Malvern Zetasizer Ultra Red. Encapsulation was determined by Ribogreen nucleic acid quantification assay (Invitrogen, R11490)

### Cell culture and virus

Human embryonic lung fibroblasts (MRC-5; CCL-171) and HCMV TB40/E (VR-3348), AD-169 (CR-538), and Merlin (VR-1590) were purchased from American Type Culture Collection (ATCC). MRC-5 were maintained in Dulbecco’s modified Eagle medium (DMEM, Gibco, 11995-065), supplemented with 10% heat-inactivated fetal bovine serum (HI-FBS, Gibco, 12484-028) and 1% penicillin/streptomycin (Gibco, 15140122). HCMV stocks were propagated by MRC-5 infection for 21 days, titered by TCID_50_ assay, and stored in 20% HI-FBS at −80 °C.

### HCMV infection assay

MRC-5s were seeded at a density of 50,000 cells per well in a 24-well plate and left overnight to allow adherence. The next day, cells were infected with HCMV at an MOI of 0.5 for 2 hours with gentle shaking every 30 minutes. Virus-containing media were then removed and replaced with antibiotic-free, supplemented DMEM. For prophylactic treatment, cells received LNP treatment 6 hours prior to infection. For therapeutic treatments, cells were treated immediately after infection. For the GCV benchmarking study, a stock GCV (G2536, Sigma-Aldrich) solution was prepared in pH 7.2 PBS (Gibco, 20012-027), diluted to the appropriate concentration and added to HCMV-infected cells.

### Flow cytometry quantification of infection

Infected cells were lifted using TrypLE Express (Gibco, 1260528), washed once with pH 7.2 PBS (Gibco, 20012-027) and stained with LIVE/DEAD™ Near-IR dead cell stain (Invitrogen, L10119) at a 1:500 dilution in pH 7.2 PBS for 30 minutes. Cells were then washed twice with cold flow cytometry staining buffer (1% FBS and 0.1% Sodium Azide in pH 7.2 PBS) before fixing in 1% paraformaldehyde (PFA, Thermo Scientific, 043368.9M) for 10 minutes. Cells were then washed twice with cold flow cytometry staining buffer to remove PFA and stored in cold flow cytometry staining buffer at 4 °C until flow cytometry analysis. Flow cytometry was performed on a Becton Dickinson Bioscience LSRFortessa X-20 analytical flow cytometer. Excitation wavelengths and emission collection windows were, respectively, GFP (488 nm, 525/50 nm) and viability dye (639 nm, 780/60 nm). All analyses of FCS files were performed using FlowJo Version 10.

### Quantification of HCMV DNA load

At the experiment endpoint, the supernatant of infected cells was removed, lifted with TrypLE Express, and resuspended in RNALater (Invitrogen, AM7021) to preserve nucleic acid integrity. Total DNA was extracted using Monarch spin gDNA extraction kit (New England Biolabs, T3010) as per manufacturer instructions, eluted in nuclease-free water and stored in −20°C until qPCR analysis. 20 ul qPCR reactions were prepared according to Genesig CMV genome quantification kit (Primer Design, R00400) with 5ng of total DNA and Low ROX PrecisionPLUS qPCR master mix (Primer Design, Z-PPLUS-LR). qPCR was performed in QuantStudio 3 real-time PCR system with the following cycling condition: 95°C for 2 mins, 50 cycles of 95°C for 10 s, 60°C for 1 min. To determine absolute viral DNA number, the standard curve method was used. To determine the relative viral load, human beta-actin (ACTB) was used as endogenous control and analyzed using the Delta Delta Ct method. 2 technical replicates were performed for each sample.

### CRISPR/Cas9 mutagenesis detection

At experiment endpoint, DNA samples were prepared as mentioned above and stored in −20°C until analysis. CRISPR/Cas9 targeted genomic and viral DNA region were amplified with Q5 high-fidelity DNA polymerase (New England Biolabs, M0491) in a PCR reaction with primers flanking the targeted/cut-anticipated region. The PCR product was then purified with Monarch Spin PCR & DNA cleanup kit (New England Biolabs, T1130) and eluted in nuclease-free water. Sanger sequencing on the amplicon was performed using the Applied Biosystem 3730xl. Tracking Indels by DEcomposition (TIDE) was used to quantify mutation frequency.

### LNP apparent pK_a_

The TNS method was used to measure the apparent pK_a_ of LNP (Cayman Chemical, 702680). Different LNPs carrying mCas9:sgAAVS1(10:1, wt/wt) were diluted with 8% sucrose, 20 mM Tris-HCl to a final lipid concentration of 500 μM, based on Ribogreen-determined RNA concentration. Buffers ranging from pH 4.0 to pH 9.0 and 83 μM TNS probe working solution were prepared as per the manufacturer’s instructions and made fresh each time. 5 μL of 500 μM LNP were added to 90 μL of the respective pH buffers, followed by 5 μL working TNS probe solution in a black 96-well plate. The mixtures were incubated for 20 minutes at room temperature on an orbital shaker. After, fluorescent levels were detected using the Tecan Infinite® 200 Pro plate reader. The pK_a_ of different LNPs were determined by fitting a three-parameter logistic equation.

## Supplementary Materials

Supplementary Data 1 and 2

Supplementary Tables S1 to S6

Supplementary Figures S1 to S13

## Acknowledgements

YMAL thanks the Ontario Graduate Scholarship, Barbara and Frank Milligan Graduate Fellowship and Emergence & Pandemic Infections Consortium Doctoral Fellowship. GT thanks the Ontario Graduate Scholarship and Precision Medicine (PRiME) Fellowship program. JP thanks the CIHR CGS-M, NSERC CGS-D and Ontario Graduate Scholarship programs. JCS thanks the Wildcat Fellows Program, Barbara and Frank Milligan Graduate Fellowship, NSERC CGS-M program, and Emergence & Pandemic Infections Consortium Doctoral Fellowship.

## Funding

University of Toronto’s Medicine by Design initiative and Pivotal Experiment Fund Round 3 which received funding from the Canada First Research Excellence Fund (CFREF-2020)

New Frontiers in Research Fund – Exploration (NFRFE–2022–00691) Canada Research Chairs Program (CRC-2020-00097)

Natural Sciences and Engineering Research Council of Canada (RGPIN-2021-03421 and DGECR-2021-00467)

Canadian Institutes of Health Research (PTT-184009)

Canada Foundation for Innovation John R. Evans Leaders Fund and Ontario Research Fund (41323).

## Author contributions

Conceptualization: OFK, YMAL

Methodology: YMAL, OFK, VHF, AH

Investigation: YMAL, JP, MQJ, YS, KLMK, GT, AMO, JCS, AMM, RM

Visualization: YMAL

Funding acquisition: OFK

Supervision: OFK

Writing – original draft: YMAL, OFK

Writing – review & editing: YMAL, OFK

## Competing interests

OFK, YMAL, MQJ and the University of Toronto have filed an invention disclosure related to this work.

## Data and material availability

All data needed to evaluate the conclusions in the paper are present in the paper or the Supplementary Materials.

## References

1. Cannon, M. J., Schmid, D. S. & Hyde, T. B. Review of cytomegalovirus seroprevalence and demographic characteristics associated with infection. Rev. Med. Virol. 20, 202–213 (2010).

2. Griffiths, P. & Reeves, M. Pathogenesis of human cytomegalovirus in the immunocompromised host. Nat. Rev. Microbiol. 19, 759–773 (2021).

3. Deayton, J. R. et al. Importance of cytomegalovirus viraemia in risk of disease progression and death in HIV-infected patients receiving highly active antiretroviral therapy. The Lancet 363, 2116–2121 (2004).

4. Haidar, G., Boeckh, M. & Singh, N. Cytomegalovirus Infection in Solid Organ and Hematopoietic Cell Transplantation: State of the Evidence. J. Infect. Dis. 221, S23–S31 (2020).

5. Swanson, M. R., Haisley, L. D., Dobyns, W. B. & Schleiss, M. R. Beyond hearing loss: exploring neurological and neurodevelopmental sequelae in asymptomatic congenital cytomegalovirus infection. Pediatr. Res. 1–12 (2025) doi:10.1038/s41390-025-04232-5.

6. Keyvanfar, A. et al. Cytomegalovirus infection contributes to acute rejection in solid organ transplant recipients: a systematic review and meta-analysis. BMC Infect. Dis. 25, 1205 (2025).

7. Kempen, J. H. et al. Mortality Risk for Patients with Cytomegalovirus Retinitis and Acquired Immune Deficiency Syndrome. Clin. Infect. Dis. 37, 1365–1373 (2003).

8. Fang, M. et al. Gastrointestinal cytomegalovirus infection in persons with HIV: a retrospective case series study. BMC Infect. Dis. 25, 506 (2025).

9. Frederick, A. W., Kitchell, E., McCormick-Baw, C., Kukkar, V. & Jain, M. K. Persistent CMV pneumonitis in HIV infection: a case report. BMC Infect. Dis. 23, 842 (2023).

10. Poh, K. C. & Zheng, S. A rare case of CMV pneumonia in HIV-infection. Respir. Med. Case Rep. 28, 100945 (2019).

11. Kawegere, E. & Goldberg, T. Atypical presentation and diagnosis of AIDS-related CMV encephalitis. BMJ Case Rep. 15, e249902 (2022).

12. Patel, R. P., Seymour, A., Brady, S. & Kahwash, R. Cytomegalovirus Encephalitis in a Heart Transplant Recipient Presenting With Behavioral Changes and Functional Neurologic Deficits. JACC Case Rep. 30, 104675 (2025).

13. Paya, C. et al. Efficacy and Safety of Valganciclovir vs. Oral Ganciclovir for Prevention of Cytomegalovirus Disease in Solid Organ Transplant Recipients. Am. J. Transplant. 4, 611–620 (2004).

14. Putera, I. et al. Antiviral therapy for cytomegalovirus retinitis: A systematic review and metaanalysis. Surv. Ophthalmol. 70, 215–231 (2025).

15. Shim, G. H. Treatment of congenital cytomegalovirus infection. Clin. Exp. Pediatr. 66, 384–394 (2022).

16. Takahata, M. et al. Occurrence of adverse events caused by valganciclovir as pre-emptive therapy for cytomegalovirus infection after allogeneic stem cell transplantation is reduced by low-dose administration. Transpl. Infect. Dis. 17, 810–815 (2015).

17. Guo, D. et al. Efficacy and toxicity analysis of ganciclovir in patients with cytomegalovirus lung infection: what is new for target range of therapeutic drug monitoring. Microbiol. Spectr. 13, e00461–25 (2025).

18. Kleiboeker, S. B. Prevalence of cytomegalovirus antiviral drug resistance in transplant recipients. Antiviral Res. 215, 105623 (2023).

19. Chou, S. et al. Comparative Emergence of Maribavir and Ganciclovir Resistance in a Randomized Phase 3 Clinical Trial for Treatment of Cytomegalovirus Infection. J. Infect. Dis. 231, e470–e477 (2025).

20. Aguado, J. M., Navarro, D., Montoto, C., Yébenes, M. & de Castro-Orós, I. Incidence of refractory CMV infection with or without antiviral resistance in Spain: A systematic literature review. Transplant. Rev. 38, 100804 (2024).

21. Avery, R. K. et al. Maribavir for Refractory Cytomegalovirus Infections With or Without Resistance Post-Transplant: Results From a Phase 3 Randomized Clinical Trial. Clin. Infect. Dis. 75, 690–701 (2022).

22. Avery, R. K. et al. Outcomes in Transplant Recipients Treated with Foscarnet for Ganciclovir-Resistant or Refractory Cytomegalovirus Infection. Transplantation 100, e74–e80 (2016).

23. Chou, S. et al. Drug Resistance Assessed in a Phase 3 Clinical Trial of Maribavir Therapy for Refractory or Resistant Cytomegalovirus Infection in Transplant Recipients. J. Infect. Dis. 229, 413–421 (2024).

24. Xiao, J. et al. Targeting human cytomegalovirus IE genes by CRISPR/Cas9 nuclease effectively inhibits viral replication and reactivation. Arch. Virol. 165, 1827–1835 (2020).

25. Gergen, J. et al. Multiplex CRISPR/Cas9 system impairs HCMV replication by excising an essential viral gene. PloS One 13, e0192602 (2018).

26. van Diemen, F. R. et al. CRISPR/Cas9-Mediated Genome Editing of Herpesviruses Limits Productive and Latent Infections. PLoS Pathog. 12, e1005701 (2016).

27. Pardi, N. et al. Expression kinetics of nucleoside-modified mRNA delivered in lipid nanoparticles to mice by various routes. J. Controlled Release 217, 345–351 (2015).

28. Hassett, K. J. et al. Optimization of Lipid Nanoparticles for Intramuscular Administration of mRNA Vaccines. Mol. Ther. Nucleic Acids 15, 1 (2019).

29. Sahin, U. et al. BNT162b2 vaccine induces neutralizing antibodies and poly-specific T cells in humans. Nature 595, 572–577 (2021).

30. Coelho, T. et al. Safety and efficacy of RNAi therapy for transthyretin amyloidosis. N. Engl. J. Med. 369, 819–829 (2013).

31. Humar, A., Siegal, D., Moussa, G. & Kumar, D. A Prospective Assessment of Valganciclovir for the Treatment of Cytomegalovirus Infection and Disease in Transplant Recipients. J. Infect. Dis. 192, 1154–1157 (2005).

32. Mattes, F. M. et al. Kinetics of Cytomegalovirus Load Decrease in Solid-Organ Transplant Recipients after Preemptive Therapy with Valganciclovir. J. Infect. Dis. 191, 89–92 (2005).

33. Perrottet, N. et al. Variable viral clearance despite adequate ganciclovir plasma levels during valganciclovir treatment for cytomegalovirus disease in D+/R-transplant recipients. BMC Infect. Dis. 10, 2 (2010).

34. Cudini, J. et al. Human cytomegalovirus haplotype reconstruction reveals high diversity due to superinfection and evidence of within-host recombination. Proc. Natl. Acad. Sci. U. S. A. 116, 5693–5698 (2019).

35. Renzette, N., Bhattacharjee, B., Jensen, J. D., Gibson, L. & Kowalik, T. F. Extensive Genome-Wide Variability of Human Cytomegalovirus in Congenitally Infected Infants. PLOS Pathog. 7, e1001344 (2011).

36. Dunn, W. et al. Functional profiling of a human cytomegalovirus genome. Proc. Natl. Acad. Sci. 100, 14223–14228 (2003).

37. Zeltzer, S. et al. Virus Control of Trafficking from Sorting Endosomes. mBio 9, e00683–18 (2018).

38. Karleuša, L., Mahmutefendić, H., Tomaš, M. I., Zagorac, G. B. & Lučin, P. Landmarks of endosomal remodeling in the early phase of cytomegalovirus infection. Virology 515, 108–122 (2018).

39. Bogdanow, B., Phan, Q. V. & Wiebusch, L. Emerging Mechanisms of G1/S Cell Cycle Control by Human and Mouse Cytomegaloviruses. mBio 12, e02934–21.

40. Alwine, J. C. The Human Cytomegalovirus Assembly Compartment: A Masterpiece of Viral Manipulation of Cellular Processes That Facilitates Assembly and Egress. PLoS Pathog. 8, e1002878 (2012).

41. Paramasivam, P. et al. Endosomal escape of delivered mRNA from endosomal recycling tubules visualized at the nanoscale. J. Cell Biol. 221, e202110137 (2021).

42. Tirosh, O. et al. The Transcription and Translation Landscapes during Human Cytomegalovirus Infection Reveal Novel Host-Pathogen Interactions. PLOS Pathog. 11, e1005288 (2015).

43. Brinkman, E. K., Chen, T., Amendola, M. & van Steensel, B. Easy quantitative assessment of genome editing by sequence trace decomposition. Nucleic Acids Res. 42, e168 (2014).

44. Tilstra, G. et al. Iterative Design of Ionizable Lipids for Intramuscular mRNA Delivery. J. Am. Chem. Soc. 145, 2294–2304 (2023).

45. Couture-Senécal, J., Tilstra, G. & Khan, O. F. Engineering ionizable lipids for rapid biodegradation balances mRNA vaccine efficacy and tolerability. 2024.08.02.606386 Preprint at 10.1101/2024.08.02.606386 (2024).

46. Patel, P., Ibrahim, N. M. & Cheng, K. The Importance of Apparent pKa in the Development of Nanoparticles Encapsulating siRNA and mRNA. Trends Pharmacol. Sci. 42, 448–460 (2021).

47. Pari, G. S. Nuts and Bolts of Human Cytomegalovirus Lytic DNA Replication. in Human Cytomegalovirus (eds Shenk, T. E. & Stinski, M. F.) 153–166 (Springer, Berlin, Heidelberg, 2008). doi:10.1007/978-3-540-77349-8_9.

48. McSharry, J. J. et al. Rapid Ganciclovir Susceptibility Assay Using Flow Cytometry for Human Cytomegalovirus Clinical Isolates. Antimicrob. Agents Chemother. 42, 2326–2331 (1998).

49. Wolf, D. G., Yaniv, I., Ashkenazi, S. & Honigman, A. Emergence of Multiple Human Cytomegalovirus Ganciclovir-Resistant Mutants with Deletions and Substitutions within the UL97 Gene in a Patient with Severe Combined Immunodeficiency. Antimicrob. Agents Chemother. 45, 593–595 (2001).

50. Walter, M., Perrone, R. & Verdin, E. Targeting Conserved Sequences Circumvents the Evolution of Resistance in a Viral Gene Drive against Human Cytomegalovirus. J. Virol. 95, 10.1128/jvi.00802-21 (2021).

51. Chen, H., Beardsley, G. P. & Coen, D. M. Mechanism of ganciclovir-induced chain termination revealed by resistant viral polymerase mutants with reduced exonuclease activity. Proc. Natl. Acad. Sci. 111, 17462–17467 (2014).

52. Stinski, M. F. & Meier, J. L. Immediate–early viral gene regulation and function. in Human Herpesviruses: Biology, Therapy, and Immunoprophylaxis (eds Arvin, A. et al.) (Cambridge University Press, Cambridge, 2007).

53. Marchini, A., Liu, H. & Zhu, H. Human cytomegalovirus with IE-2 (UL122) deleted fails to express early lytic genes. J. Virol. 75, 1870–1878 (2001).

54. Michaelis, M. et al. The multi-targeted kinase inhibitor sorafenib inhibits human cytomegalovirus replication. Cell. Mol. Life Sci. CMLS 68, 1079–1090 (2011).

55. Lim, W. Y., Lee, J. H., Choi, Y. & Yoon, K. Verteporfin is an effective inhibitor of HCMV replication. Virus Res. 350, 199475 (2024).

56. Yasar, H. et al. Kinetics of mRNA delivery and protein translation in dendritic cells using lipid-coated PLGA nanoparticles. J. Nanobiotechnology 16, 72 (2018).

57. Leonhardt, C. et al. Single-cell mRNA transfection studies: Delivery, kinetics and statistics by numbers. Nanomedicine Nanotechnol. Biol. Med. 10, 679–688 (2014).

58. Chou, S. Cytomegalovirus UL97 mutations in the era of ganciclovir and maribavir. Rev. Med. Virol. 18, 233–246 (2008).

59. Gentry, B. G. & Drach, J. C. Metabolism of Cyclopropavir and Ganciclovir in Human Cytomegalovirus-Infected Cells. Antimicrob. Agents Chemother. 58, 2329–2333 (2014).

60. Li, T. et al. CRISPR/Cas9 therapeutics: progress and prospects. Signal Transduct. Target. Ther. 8, 36 (2023).

61. Chang, H. H. Y., Pannunzio, N. R., Adachi, N. & Lieber, M. R. Non-homologous DNA end joining and alternative pathways to double-strand break repair. Nat. Rev. Mol. Cell Biol. 18, 495–506 (2017).

62. Luo, M. H., Rosenke, K., Czornak, K. & Fortunato, E. A. Human Cytomegalovirus Disrupts both Ataxia Telangiectasia Mutated Protein (ATM)- and ATM-Rad3-Related Kinase-Mediated DNA Damage Responses during Lytic Infection. J. Virol. 81, 1934–1950 (2007).

63. E, X. & Kowalik, T. F. The DNA Damage Response Induced by Infection with Human Cytomegalovirus and Other Viruses. Viruses 6, 2155–2185 (2014).

64. Wu, X., Zhou, X., Wang, S. & Mao, G. DNA damage response(DDR): a link between cellular senescence and human cytomegalovirus. Virol. J. 20, 250 (2023).

65. Marraffini, L. A. & Sontheimer, E. J. CRISPR Interference Limits Horizontal Gene Transfer in Staphylococci by Targeting DNA. Science 322, 1843–1845 (2008).

