## Supplementary Information for "The Clearance of Human Cytomegalovirus Using CRISPR/Cas9 RNA Lipid Nanoparticles"

### 1. Supplementary Tables

**Supplementary Table S1: Dispensability analysis of viral gene the selected GO: biological processes**

| GO: Biological Processes | ORFs | In vitro Growth (Fibroblasts) | Growth condition |
|---|---|---|---|
| Mitigation of host antiviral defense response (GO:0019049) | US2 | Dispensable[1] | Wild-type growth[1] |
| | US3 | Dispensable[1] | Wild-type growth[1] |
| | US7 | Dispensable[1] | Wild-type growth[1] |
| | US8 | Dispensable[1] | Wild-type growth[1] |
| | US9 | Dispensable[1] | Wild-type growth[1] |
| | US10 | Dispensable[1] | Wild-type growth[1] |
| | US11 | Dispensable[1] | Wild-type growth[1] |
| | US27 | Dispensable[1] | Wild-type growth[1] |
| | US28 | Dispensable[1] | Wild-type growth[1] |
| | US33 | Dispensable[1] | Wild-type growth[1] |
| | UL146 | Dispensable[1] | Wild-type growth[1] |
| | UL119/UL118 | Dispensable[1] | Wild-type growth[1] |
| Bidirectional double-stranded viral DNA replication (GO:0039686) | UL44 | Essential[1] | No Growth[1] |
| | UL54 | Essential[1] | No Growth[1] |
| | UL57 | Essential[1] | No Growth[1] |
| | UL70 | Essential[1] | No Growth[1] |
| | UL102 | Essential[1] | No Growth[1] |
| | UL105 | Essential[1] | No Growth[1] |
| | UL112/113 | Essential[1] | Severe growth defects[1] |

| | | | |
|---|---|---|---|
| | UL122 | Essential[1] | No Growth[1] |
| | UL123 | Essential[1] | No Growth[1] |
| | IRS1 | Dispensable[1] | Wild-type growth[1] |
| Viral release from host cell (GO:0019076) | UL51 | Essential[1] | No Growth[1] |
| | UL56 | Essential[1] | No Growth[1] |
| | UL77 | Essential[1] | No Growth[1] |
| | UL80 | Essential[1] | No Growth[1] |
| | UL89 | Essential[4] | Severe growth defects[4] |
| | UL93 | Essential[1] | No Growth[1] |
| | UL104 | Essential[1] | No Growth |
| Viral entry into host cell (GO:0046718) | UL55 | Essential[1] | No Growth |
| | UL75 | Essential[1] | No Growth |
| | UL77 | Essential[1] | No Growth |
| | UL115 | Essential[1] | No Growth |
| | UL128 | Dispensable[5] (in fibroblast) | Enhanced growth[5] (Knock-in was detrimental to growth). Essential for endothelial and endothelial cells infection[5] |
| | UL130 | Dispensable[5] (in fibroblast) | Enhanced growth[5] (Knock-in was detrimental to growth). Essential for endothelial and endothelial cells infection[5] |

| | | | |
|---|---|---|---|
| | UL131 | Dispensable[5] (in fibroblast) | Enhanced growth[5] (Knock-in was detrimental to growth).<br><br>Essential for endothelial and endothelial cells infection[5] |
| Suppression by virus of host type I interferon-mediated signaling pathway (GO:0039502) | UL23 | Dispensable | Enhanced growth[1] |
| | UL111A | Dispensable[1] | Wild-type growth |
| | UL123 | Essential[1] | Severe growth defects[1] |
| | UL145 | Dispensable[6] | Wild-type growth[6] |
| | TRS1 | Dispensable[1,7] | Minor growth defects If IRS1 is expressed[1,7] |
| | IRS1 | Dispensable[1,7] | Minor growth defects If TRS1 is expressed[1,7] |

**Supplementary Table S2: Cas9 mRNA coding sequence**

| Cas9 mRNA coding sequence (5’ to 3’) |
|---|
| TAATACGACTCACTATAAGGGAGACCCAAGCTGGCTAGCCGCCACCATGGCCCCC<br>AAGAAGAAGCGGAAGGTGGGCATCCACGGCGTGCCCGCCGCCGACAAGAAGTAC<br>AGCATCGGCCTGGACATCGGCACCAACAGCGTGGGCTGGGCCGTGATCACCGACG<br>AGTACAAGGTGCCCAGCAAGAAGTTCAAGGTGCTGGGCAACACCGACCGGCACA<br>GCATCAAGAAGAACCTGATCGGCGCCCTGCTGTTCGACAGCGGCGAGACCGCCGA<br>GGCCACCCGGCTGAAGCGGACCGCCCGGCGGCGGTACACCCGGCGGAAGAACCG<br>GATCTGCTACCTGCAGGAGATCTTCAGCAACGAGATGGCCAAGGTGGACGACAGC<br>TTCTTCCACCGGCTGGAGGAGAGCTTCCTGGTGGAGGAGGACAAGAAGCACGAGC<br>GGCACCCCATCTTCGGCAACATCGTGGACGAGGTGGCCTACCACGAGAAGTACCC<br>CACCATCTACCACCTGCGGAAGAAGCTGGTGGACAGCACCGACAAGGCCGACCTG<br>CGGCTGATCTACCTGGCCCTGGCCCACATGATCAAGTTCCGGGGCCACTTCCTGAT<br>CGAGGGCGACCTGAACCCCGACAACAGCGACGTGGACAAGCTGTTCATCCAGCTG<br>GTGCAGACCTACAACCAGCTGTTCGAGGAGAACCCCATCAACGCCAGCGGCGTGG<br>ACGCCAAGGCCATCCTGAGCGCCCGGCTGAGCAAGAGCCGGCGGCTGGAGAACC<br>TGATCGCCCAGCTGCCCGGCGAGAAGAAGAACGGCCTGTTCGGCAACCTGATCGC<br>CCTGAGCCTGGGCCTGACCCCCAACTTCAAGAGCAACTTCGACCTGGCCGAGGAC<br>GCCAAGCTGCAGCTGAGCAAGGACACCTACGACGACGACCTGGACAACCTGCTGG<br>CCCAGATCGGCGACCAGTACGCCGACCTGTTCCTGGCCGCCAAGAACCTGAGCGA<br>CGCCATCCTGCTGAGCGACATCCTGCGGGTGAACACCGAGATCACCAAGGCCCCC<br>CTGAGCGCCAGCATGATCAAGCGGTACGACGAGCACCACCAGGACCTGACCCTGC<br>TGAAGGCCCTGGTGCGGCAGCAGCTGCCCGAGAAGTACAAGGAGATCTTCTTCGA<br>CCAGAGCAAGAACGGCTACGCCGGCTACATCGACGGCGGCGCCAGCCAGGAGGA<br>GTTCTACAAGTTCATCAAGCCCATCCTGGAGAAGATGGACGGCACCGAGGAGCTG<br>CTGGTGAAGCTGAACCGGGAGGACCTGCTGCGGAAGCAGCGGACCTTCGACAAC<br>GGCAGCATCCCCCACCAGATCCACCTGGGCGAGCTGCACGCCATCCTGCGGCGGC<br>AGGAGGACTTCTACCCCTTCCTGAAGGACAACCGGGAGAAGATCGAGAAGATCCT<br>GACCTTCCGGATCCCCTACTACGTGGGCCCCCTGGCCCGGGGCAACAGCCGGTTC<br>GCCTGGATGACCCGGAAGAGCGAGGAGACCATCACCCCCTGGAACTTCGAGGAG<br>GTGGTGGACAAGGGCGCCAGCGCCCAGAGCTTCATCGAGCGGATGACCAACTTCG<br>ACAAGAACCTGCCCAACGAGAAGGTGCTGCCCAAGCACAGCCTGCTGTACGAGTA<br>CTTCACCGTGTACAACGAGCTGACCAAGGTGAAGTACGTGACCGAGGGCATGCGG<br>AAGCCCGCCTTCCTGAGCGGCGAGCAGAAGAAGGCCATCGTGGACCTGCTGTTCA<br>AGACCAACCGGAAGGTGACCGTGAAGCAGCTGAAGGAGGACTACTTCAAGAAGA<br>TCGAGTGCTTCGACAGCGTGGAGATCAGCGGCGTGGAGGACCGGTTCAACGCCAG<br>CCTGGGCACCTACCACGACCTGCTGAAGATCATCAAGGACAAGGACTTCCTGGAC<br>AACGAGGAGAACGAGGACATCCTGGAGGACATCGTGCTGACCCTGACCCTGTTCG<br>AGGACCGGGAGATGATCGAGGAGCGGCTGAAGACCTACGCCCACCTGTTCGACG<br>ACAAGGTGATGAAGCAGCTGAAGCGGCGGCGGTACACCGGCTGGGGCCGGCTGA<br>GCCGGAAGCTGATCAACGGCATCCGGGACAAGCAGAGCGGCAAGACCATCCTGG<br>ACTTCCTGAAGAGCGACGGCTTCGCCAACCGGAACTTCATGCAGCTGATCCACGA<br>CGACAGCCTGACCTTCAAGGAGGACATCCAGAAGGCCCAGGTGAGCGGCCAGGG<br>CGACAGCCTGCACGAGCACATCGCCAACCTGGCCGGCAGCCCCGCCATCAAGAAG<br>GGCATCCTGCAGACCGTGAAGGTGGTGGACGAGCTGGTGAAGGTGATGGGCCGG |

CACAAGCCCGAGAACATCGTGATCGAGATGGCCCGGGAGAACCAGACCACCCAG
AAGGGCCAGAAGAACAGCCGGGAGCGGATGAAGCGGATCGAGGAGGGCATCAA
GGAGCTGGGCAGCCAGATCCTGAAGGAGCACCCCGTGGAGAACACCCAGCTGCA
GAACGAGAAGCTGTACCTGTACTACCTGCAGAACGGCCGGGACATGTACGTGGAC
CAGGAGCTGGACATCAACCGGCTGAGCGACTACGACGTGGACCACATCGTGCCCC
AGAGCTTCCTGAAGGACGACAGCATCGACAACAAGGTGCTGACCCGGAGCGACA
AGAACCGGGGCAAGAGCGACAACGTGCCCAGCGAGGAGGTGGTGAAGAAGATGA
AGAACTACTGGCGGCAGCTGCTGAACGCCAAGCTGATCACCCAGCGGAAGTTCGA
CAACCTGACCAAGGCCGAGCGGGGCGGCCTGAGCGAGCTGGACAAGGCCGGCTT
CATCAAGCGGCAGCTGGTGGAGACCCGGCAGATCACCAAGCACGTGGCCCAGATC
CTGGACAGCCGGATGAACACCAAGTACGACGAGAACGACAAGCTGATCCGGGAG
GTGAAGGTGATCACCCTGAAGAGCAAGCTGGTGAGCGACTTCCGGAAGGACTTCC
AGTTCTACAAGGTGCGGGAGATCAACAACTACCACCACGCCCACGACGCCTACCT
GAACGCCGTGGTGGGCACCGCCCTGATCAAGAAGTACCCCAAGCTGGAGAGCGA
GTTCGTGTACGGCGACTACAAGGTGTACGACGTGCGGAAGATGATCGCCAAGAGC
GAGCAGGAGATCGGCAAGGCCACCGCCAAGTACTTCTTCTACAGCAACATCATGA
ACTTCTTCAAGACCGAGATCACCCTGGCCAACGGCGAGATCCGGAAGCGGCCCCT
GATCGAGACCAACGGCGAGACCGGCGAGATCGTGTGGGACAAGGGCCGGGACTT
CGCCACCGTGCGGAAGGTGCTGAGCATGCCCCAGGTGAACATCGTGAAGAAGACC
GAGGTGCAGACCGGCGGCTTCAGCAAGGAGAGCATCCTGCCCAAGCGGAACAGC
GACAAGCTGATCGCCCGGAAGAAGGACTGGGACCCCAAGAAGTACGGCGGCTTC
GACAGCCCCACCGTGGCCTACAGCGTGCTGGTGGTGGCCAAGGTGGAGAAGGGC
AAGAGCAAGAAGCTGAAGAGCGTGAAGGAGCTGCTGGGCATCACCATCATGGAG
CGGAGCAGCTTCGAGAAGAACCCCATCGACTTCCTGGAGGCCAAGGGCTACAAGG
AGGTGAAGAAGGACCTGATCATCAAGCTGCCCAAGTACAGCCTGTTCGAGCTGGA
GAACGGCCGGAAGCGGATGCTGGCCAGCGCCGGCGAGCTGCAGAAGGGCAACGA
GCTGGCCCTGCCCAGCAAGTACGTGAACTTCCTGTACCTGGCCAGCCACTACGAG
AAGCTGAAGGGCAGCCCCGAGGACAACGAGCAGAAGCAGCTGTTCGTGGAGCAG
CACAAGCACTACCTGGACGAGATCATCGAGCAGATCAGCGAGTTCAGCAAGCGG
GTGATCCTGGCCGACGCCAACCTGGACAAGGTGCTGAGCGCCTACAACAAGCACC
GGGACAAGCCCATCCGGGAGCAGGCCGAGAACATCATCCACCTGTTCACCCTGAC
CAACCTGGGCGCCCCCGCCGCCTTCAAGTACTTCGACACCACCATCGACCGGAAG
CGGTACACCAGCACCAAGGAGGTGCTGGACGCCACCCTGATCCACCAGAGCATCA
CCGGCCTGTACGAGACCCGGATCGACCTGAGCCAGCTGGGCGGCGACAGCGGCG
GCAAGCGGCCCGCCGCCACCAAGAAGGCCGGCCAGGCCAAGAAGAAGAAGGGCA
GCTACCCCTACGACGTGCCCGACTACGCCTGATAAGCTGCCTTCTGCGGGGCTTGC
CTTCTGGCCATGCCCTTCTTCTCTCCCTTGCACCTGTACCTCTTGGTCTTTGAATAA
AGCCTGAGTAGGAAGAAAAAAAAAAAAAAAAAAAAAAAAAAAAAAAAAAAAAAAAAAAA
AAAAAAAAAAAAAAAAAAAAAAAAAAAAAAAAAAAAAAAAAAAAAAAAAAAAAAAAAAA
AAAAAAAAAAAATCTAGA

**Supplementary Table S3: Oligonucleotide primers used for Tracking of Indels by Decomposition (TIDE) analysis**

| Amplicon | Forward Primer (5' to 3') | Reverse Primer (5' to 3') | Size (bp) |
|---|---|---|---|
| AAVS1 | CCCCGTTCTCCTGTGGATTC | ATCCTCTCTGGCTCCATCGT | 495 |
| UL44 | GCCGAGCTGAACTCCATATTGA | CTATGACTCTGGGATGACGCC | 623 |

**Supplementary Table S4: CRISPR RNA LNP (SM-102 LNP) characteristics**

| sgRNA Target | sgRNA MW (Da) | Z-average (d.nm) | PDI | Zeta Potential (mV) | EE (%) |
|---|---|---|---|---|---|
| AAVS1 | 32511 | 97.2 | 0.116 | 6.39 | 95.2 |
| UL44 | 32454 | 96.2 | 0.119 | 8.34 | 95.7 |
| UL54 | 32360 | 93.0 | 0.113 | 10.1 | 94.7 |
| UL57 | 32370 | 95.6 | 0.143 | 13.97 | 95.2 |
| UL70 | 32392 | 95.0 | 0.157 | 7.8 | 95.7 |
| UL102 | 32330 | 127.8 | 0.304 | 12.8 | 94.9 |
| UL105 | 32346 | 95.5 | 0.107 | 12.54 | 96.1 |
| UL112/113 | 32511 | 87.8 | 0.101 | 6.1 | 95.3 |
| UL122 | 32392 | 100.8 | 0.116 | 5.54 | 94.0 |
| UL123 | 32289 | 89.3 | 0.117 | 11.0 | 95.4 |
| IRS1 | 32527 | 108.2 | 0.192 | 14.6 | 94.4 |

**Supplementary Table S5: Lipids and molar ratios used in different LNP formulations**

| Formulation | Ionizable Lipid | PEG Lipid | Molar Ratio (Ionizable Lipid: Cholesterol: DSPC: PEG Lipid) |
|---|---|---|---|
| SM-102 LNP | SM-102 | DMG-PEG 2000 | 50:38.5:10:1.5 |
| SM-102-40 LNP | SM-102 | DMG-PEG 2000 | 33:25.5:40:1.5 |
| ALC-0315 LNP | ALC-0315 | ALC-0159 | 46.3:42.7:9.4:1.6 |
| DLin-MC3-DMA LNP | DLin-MC3-DMA | PEG (2000)-C-DMG | 50:38.5:10:1.5 |
| βN2-40 LNP | βN2 | DMG-PEG 2000 | 33:25.5:40:1.5 |
| δO3-40 LNP | δO3 | DMG-PEG 2000 | 33:25.5:40:1.5 |

**Supplementary Table S6: Characterization of different LNP formulations**

| Formulation | sgRNA Target | Z-average (d.nm) | PDI | Zeta Potential (mV) | EE (%) |
|---|---|---|---|---|---|
| SM-102 LNP | AAVS1 | 97.2 | 0.116 | 6.39 | 95.2 |
| | UL44 | 96.2 | 0.119 | 8.34 | 95.7 |
| | UL57 | 95.6 | 0.143 | 13.97 | 95.2 |
| | UL105 | 95.5 | 0.107 | 12.54 | 96.1 |
| | Multiplex | 84.9 | 0.085 | 9.22 | 95.8 |
| ALC-0315 LNP | AAVS1 | 89.3 | 0.118 | -9.87 | 94.0 |
| | UL44 | 80.8 | 0.106 | -17.73 | 95.1 |
| | UL57 | 85.3 | 0.117 | -13.93 | 95.8 |
| | UL105 | 84.6 | 0.096 | -14.06 | 95.4 |
| | Multiplex | 93.1 | 0.231 | -12.94 | 95.4 |

| | | | | | |
|---|---|---|---|---|---|
| DLin-MC3-DMA LNP | AAVS1 | 131.7 | 0.170 | 1.93 | 91.3 |
| | UL44 | 125.7 | 0.203 | 1.89 | 92.4 |
| | UL57 | 138.2 | 0.182 | -1.93 | 91.5 |
| | UL105 | 135.7 | 0.1838 | 4.31 | 91.1 |
| | Multiplex | 138.1 | 0.165 | 1.48 | 91.5 |
| βN2-40 LNP | AAVS1 | 152.2 | 0.239 | 8.42 | 79.8 |
| | UL44 | 143.9 | 0.205 | 6.83 | 82.2 |
| | UL57 | 148.4 | 0.222 | 7.67 | 79.6 |
| | UL105 | 146.7 | 0.207 | 7.77 | 80.6 |
| | Multiplex | 139.4 | 0.216 | 7.98 | 84.0 |
| δO3-40 LNP | AAVS1 | 120.9 | 0.196 | 2.678 | 88.4 |
| | UL44 | 130.6 | 0.283 | 4.02 | 96.8 |
| | UL57 | 108.8 | 0.282 | 4.35 | 96.5 |
| | UL105 | 105.9 | 0.255 | 3.60 | 96.1 |
| | Multiplex | 94.5 | 0.184 | 5.66 | 97.2 |

### 2. Supplementary Figures

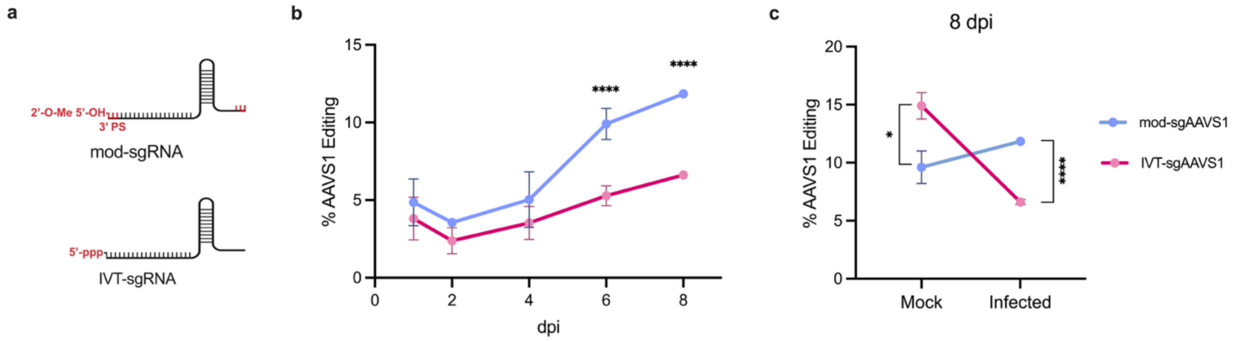


**Supplementary Fig. S1: Impact of sgRNA modification on CRISPR RNA LNP gene editing during HCMV infection.** **(a)** Diagram illustrating the main differences between modified (mod) sgRNA and in vitro transcribed (IVT) sgRNA. Mod-sgRNA contains a 2' O-methyl analog on the first and last three bases, 5' hydroxyl, and 3' phosphorothioate antinucleotide linkages between the first three and last two bases. IVT-sgRNA contains an uncapped 5' triphosphate. **(b)** Percentage AAVS1 gene editing after receiving daily treatments of CRISPR RNA LNP containing either mod-sgRNA or IVT-sgRNA at respective days post-infection (dpi) **(c)** The impact of sgRNA modification and HCMV infection on AAVS1 gene editing.

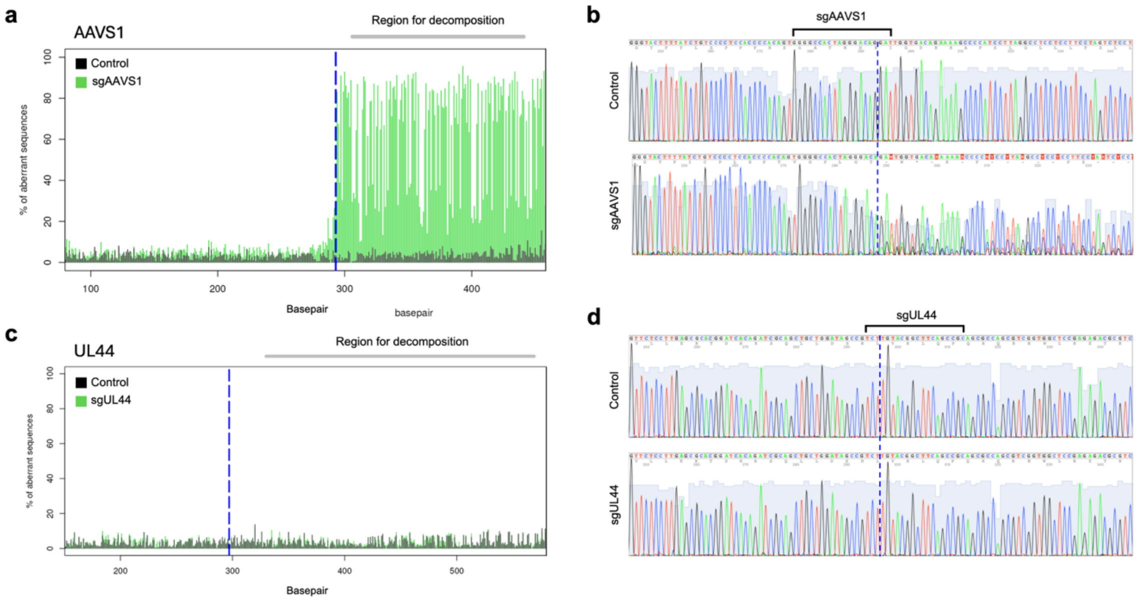


**Supplementary Fig. S2: TIDE analysis of AAVS1 and UL44 amplicon after receiving sgAAVS1 or sgUL44 CRISPR RNA LNP, respectively, during HCMV infection.** Average percentages of conflicting nucleotide (aberrant sequences) within the region for decomposition for **(a)** AAVS1 and **(c)** UL44 amplicon after receiving respective CRISPR RNA LNP treatment. Black traces represent amplicons that did not receive CRISPR/Cas9 treatment (Control). Green traces represent amplicons that received corresponding CRISPR/Cas9 treatment (sgAAVS1 or sgUL44). The dotted blue line represents the expected cut site. Representative DNA chromatogram traces of **(b)** AAVS1 and **(d)** UL44 amplicon after receiving either no CRISPR/Cas9 treatment (Control), or corresponding CRISPR/Cas9 treatment (sgAAVS1 or sgUL44). The bracket depicts the spacer complementary site. The dotted blue line represents the expected cut site.

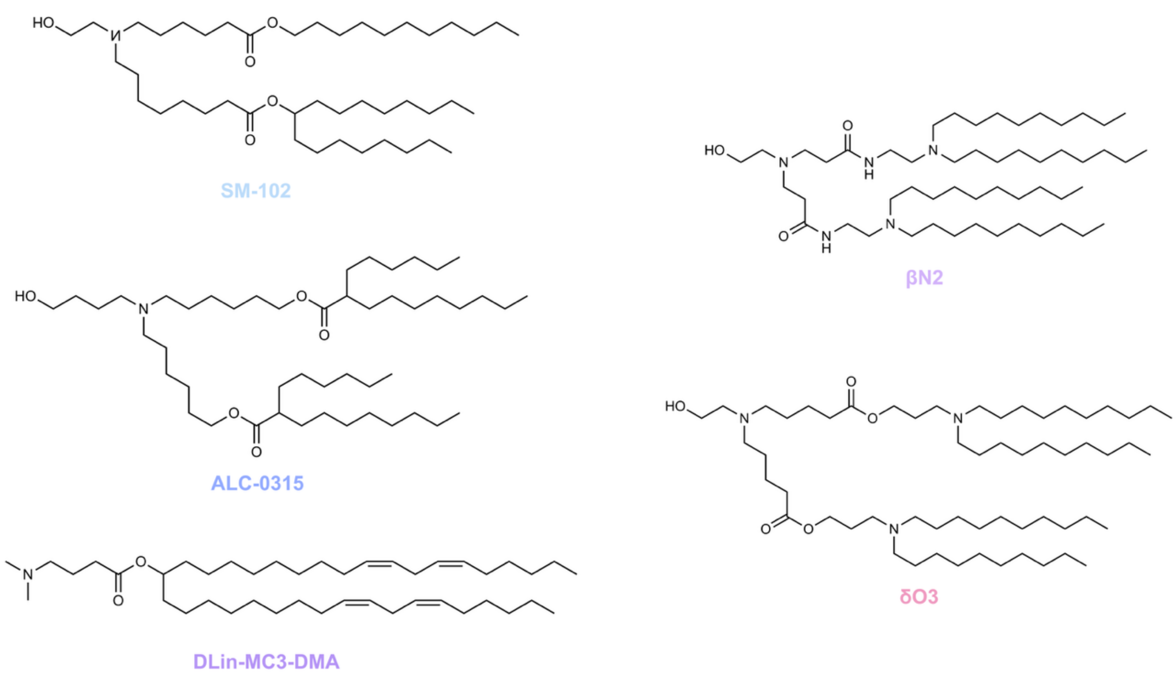


**Supplementary Figure S3: Structures of the ionizable lipids used in this study.**

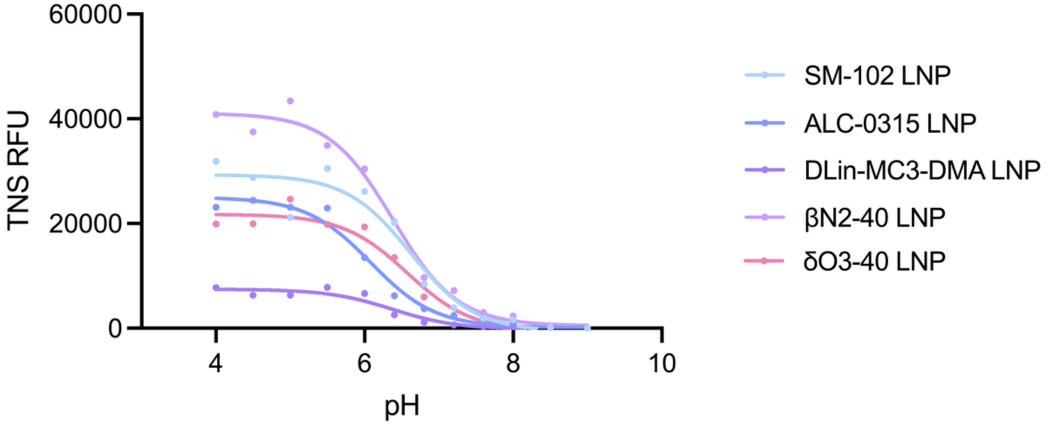


| Formulation | SM-102 LNP | ALC-0315 LNP | DLin-MC3-DMA LNP | βN2-40 LNP | δO3-40 LNP |
|---|---|---|---|---|---|
| $pK_a$ | 6.599 | 6.080 | 6.315 | 6.383 | 6.556 |

**Supplementary Fig. S4: TNS measurements of different LNP formulation and their respective $pK_a$.**

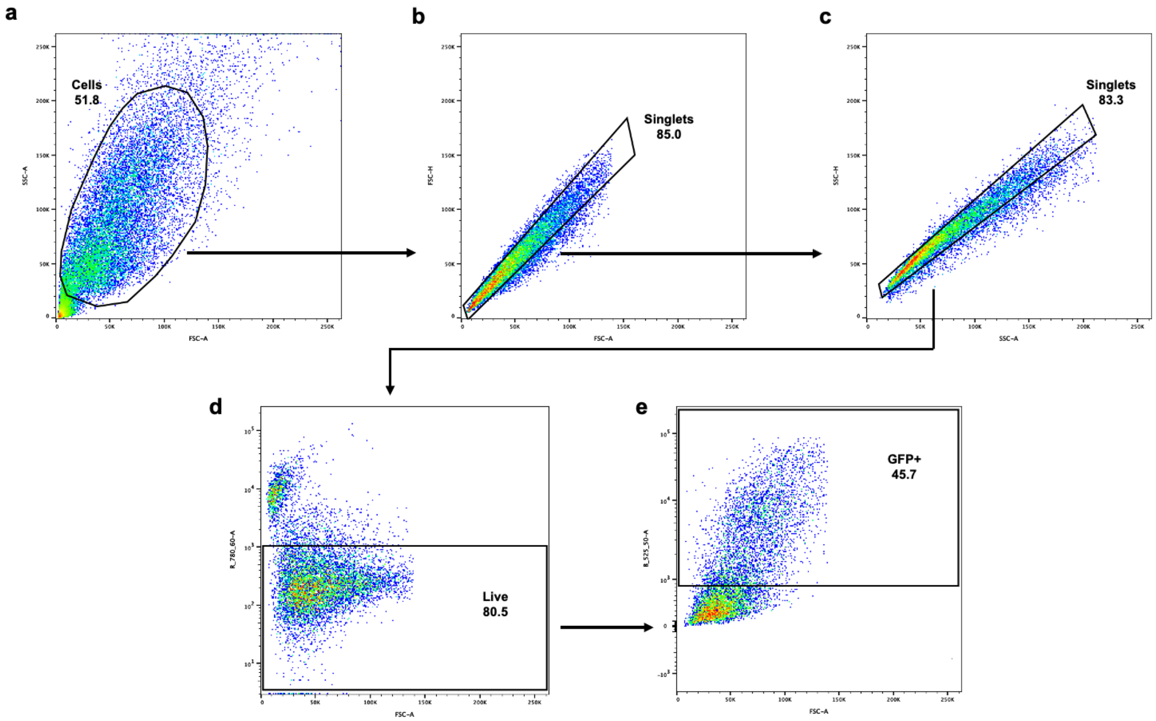


**Supplementary Fig. S5: Flow cytometry gating strategy used for evaluating TB40/E-GFP infection in MRC-5 fibroblast. (a)** Infected and non-infected MRC-5s are selected. Infected cells have shown to increase in forward (size) and side (granularity) scatter. (**b, c)** Singlets are gated to exclude doublet events. (**d)** Live cell gating. (**e)** Infected, GFP+ cells gating. Mock infection was used as a fluorescent minus one (FMO) to gate GFP+ events.

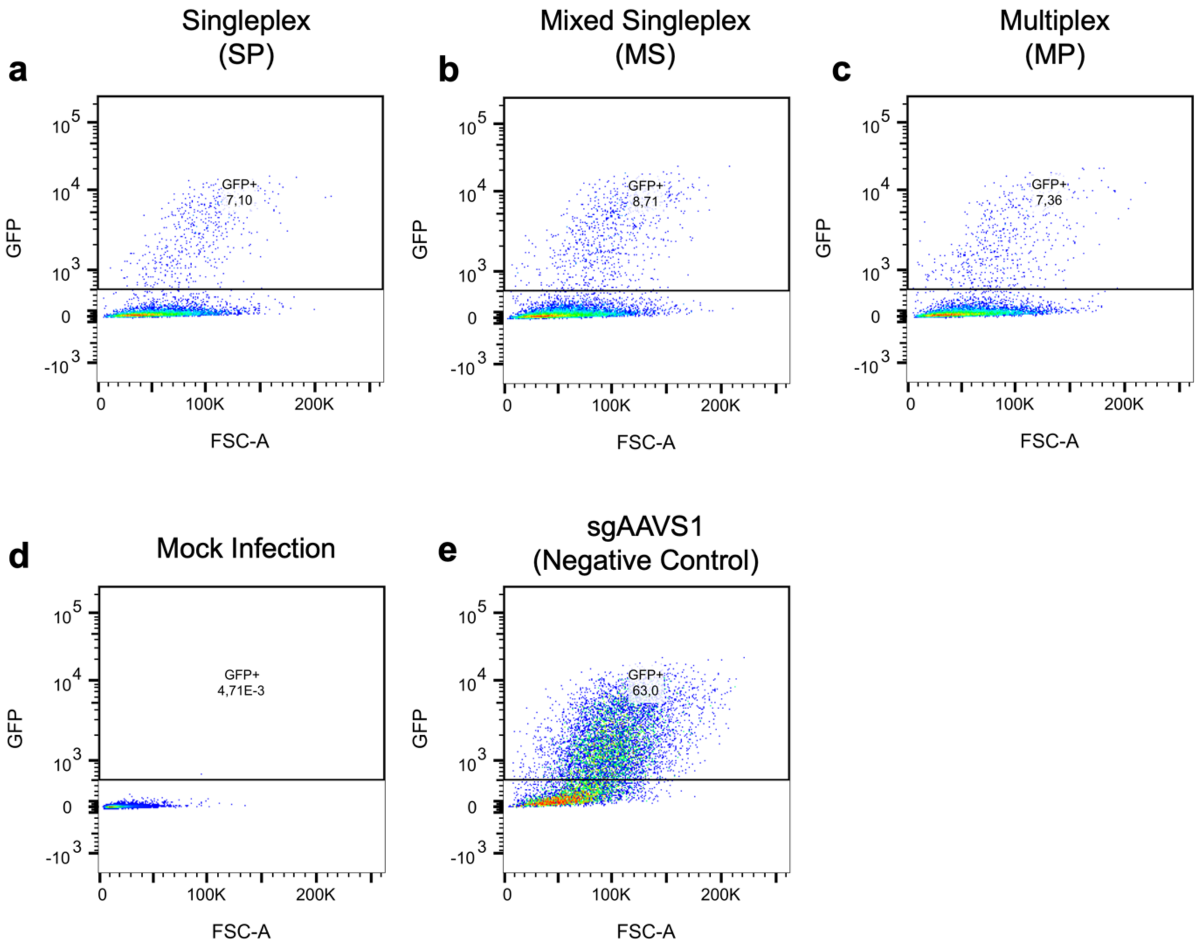


**Supplementary Fig. S6: Antiviral effects of single-dose singleplex, mixed singleplex, or multiplex βN2-40 LNP treatment.** Representative flow cytometry dot plots and GFP+ gate of quantifying HCMV infection after **(a)** singleplex, **(b)** mixed singleplex, or **(c)** multiplex βN2-40 LNP treatment. **(d)** Mock infection and **(e)** genomically targeting AAVS1 βN2-40 LNP were used as controls. The dot plot shows forward scatter area (FSC-A) versus GFP.

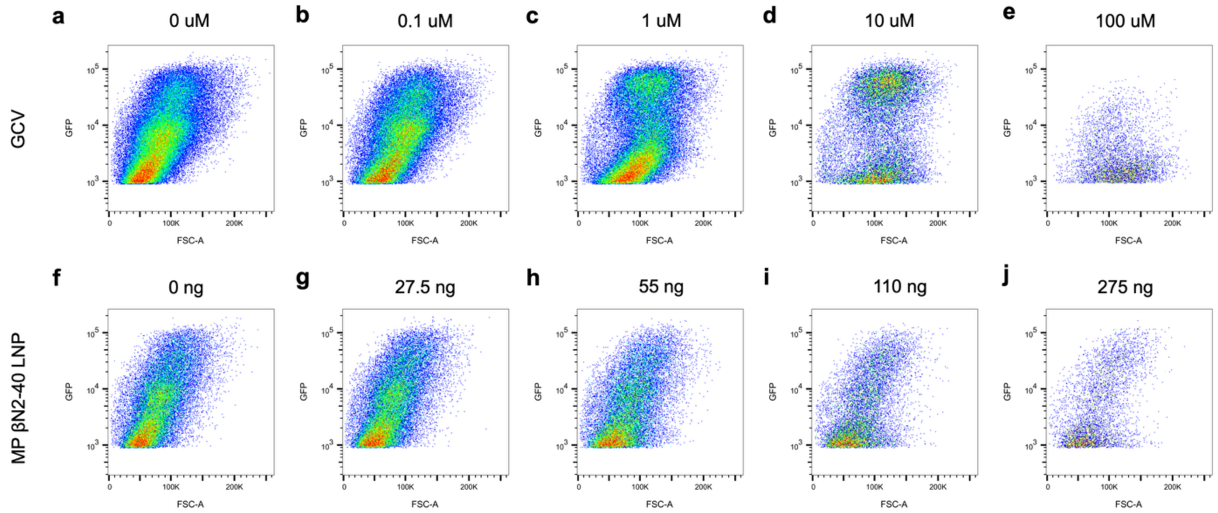


**Supplementary Fig. S7: HCMV-infected population after escalating doses of ganciclovir (GCV) or MP βN2-40 LNP treatment.** Representative flow cytometry dot plots of the infected, GFP+ population after receiving different doses of **(a-e)** GCV or **(f-j)** MP βN2-40 LNP treatment. The dot plot shows forward scatter area (FSC-A) versus GFP. The depicted doses are near and within the linear region of their respective dose-response curve. The dot plot shows FSC-A versus GFP.

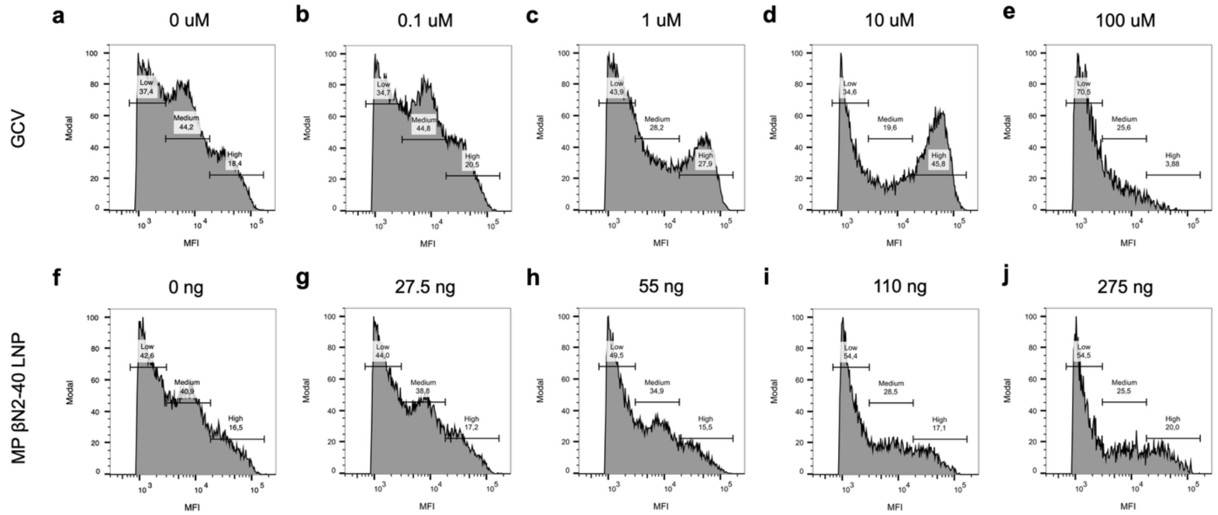


**Supplementary Fig. S8: Viral burden of HCMV-infected population after escalating doses of GCVor MP βN2-40 LNP treatment.** Representative median fluorescent intensity (MFI) histogram of the infected, GFP+ population after receiving different doses of **(a-e)** GCV or **(f-j)** MP βN2-40 LNP treatment. The depicted doses are near the linear region of their respective dose-response curves. Three distinct peaks were observed in the untreated controls and were classified as low-, medium-, and high-grade infection. The histogram shows MFI versus mode.

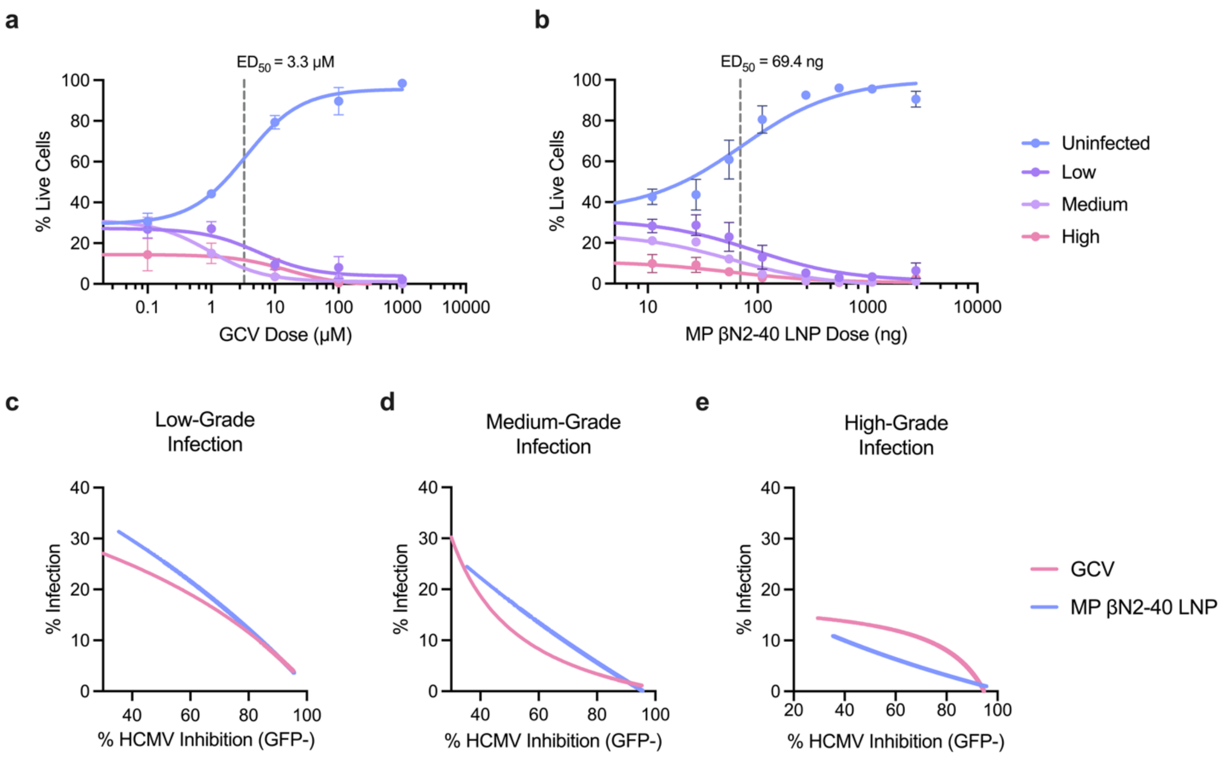


**Supplementary Fig. S9: Proportion of cells with low-, medium-, and high-grade infection after escalating doses of GCV or MP βN2-40 LNP treatment.** Percentage of uninfected cells, and cells with low-, medium-, and high-grade infection after receiving different doses of **(a)** GCV or **(b)** MP βN2-40 LNP treatment. Data points were fitted with a three-parameter sigmodal function. The dotted line depicts the median effective dose ($ED_{50}$). Antiviral kinetics of GCV (red) and MP βN2-40 LNP (blue) treatment against **(c)** low-, **(d)** medium-, and **(e)** high-grade infection at equivalent antiviral levels.

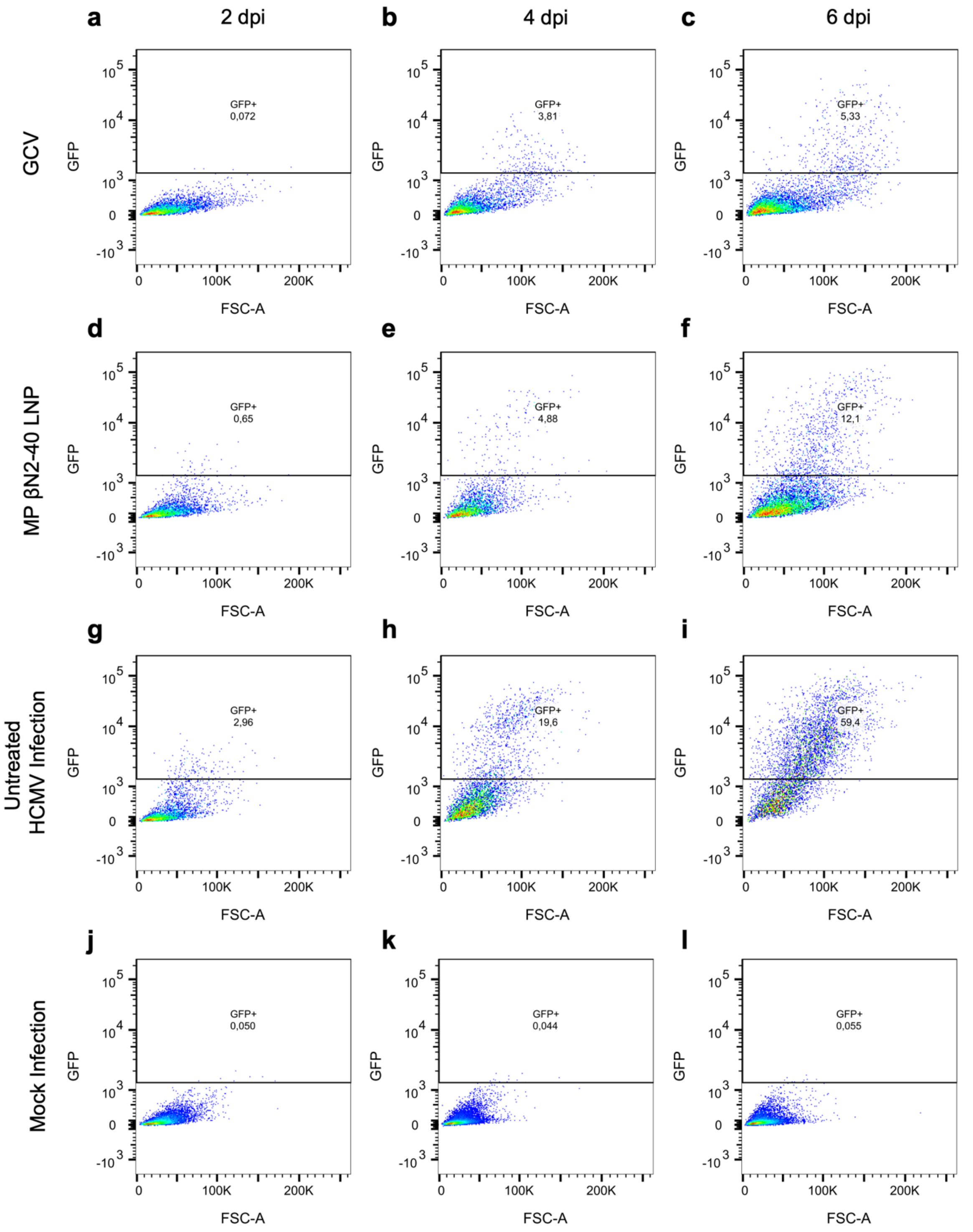


**Supplementary Fig. S10: Antiviral effect comparison between GCV and MP βN2-40 LNP treatment.** Representative flow cytometry dot plot and GFP+ gate quantifying HCMV infection after receiving a predicted equivalent single dose of **(a-c)** GCV or **(d-f)** MP βN2-40 LNP at 2,4,6 dpi. **(g-i)** Untreated HCMV infection and **(j-l)** mock infection were used as controls. The dot plot shows forward scatter area (FSC-A) versus GFP.

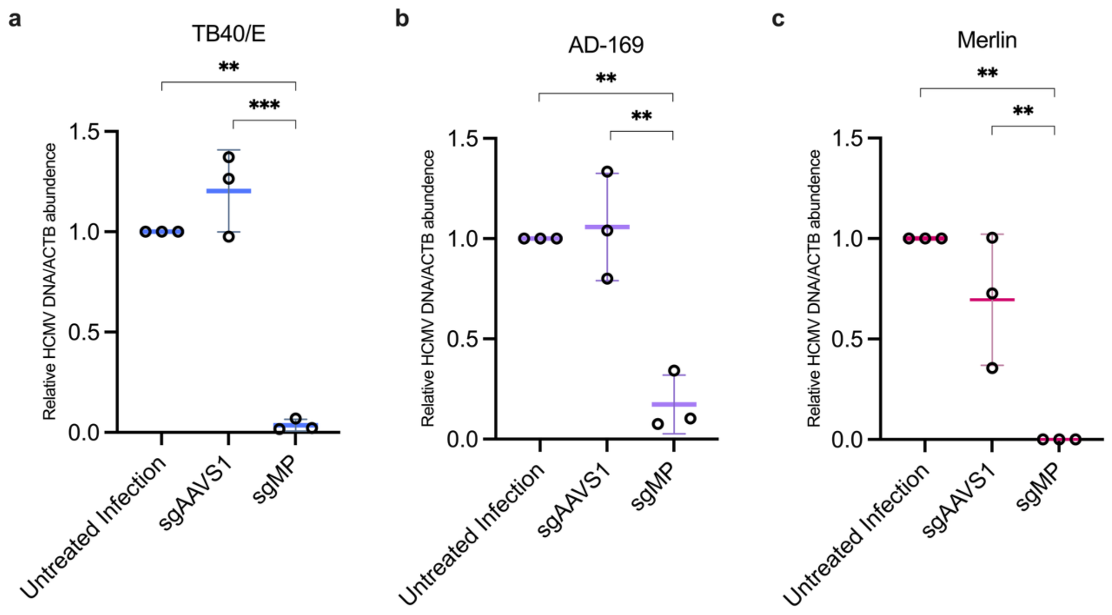


**Supplementary Fig. S11: Relative quantification of viral DNA load after MP βN2-40 LNP against different HCMV strains.** Relative quantification of (**a**) TB40/E (**b**) AD-169 and (**c**) Merlin viral DNA load after MP βN2-40 treatment. The data are shown as the mean ± standard deviation. Statistical analysis: one-way ANOVA for **a-c**. $^{*}P \leq 0.05$, $^{**}P \leq 0.01$, $^{***}P \leq 0.001$.

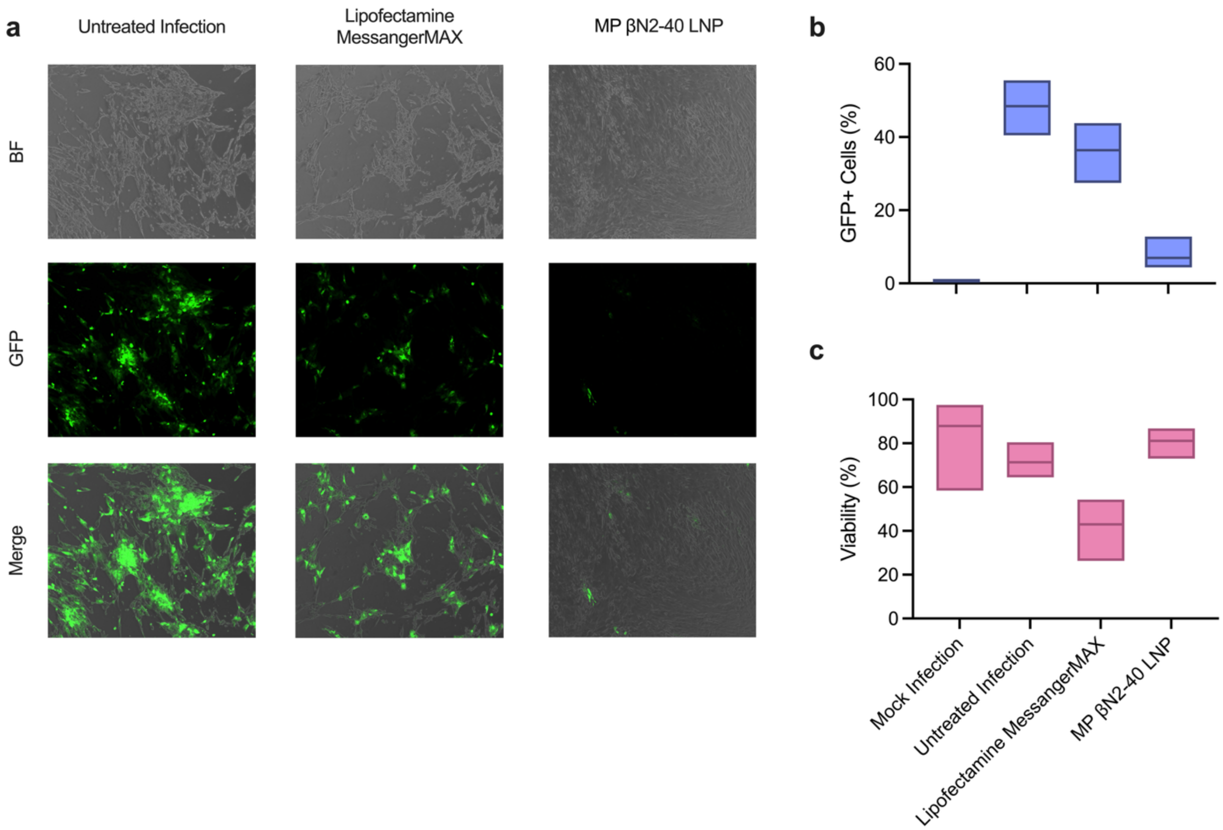


**Supplementary Fig. S12: Antiviral effects of anti-HCMV CRISPR RNA delivered by LipofectamineMessangerMax or βN2-40. (a)** Representative microscopy images of TB40/E-GFP infection after therapeutic treatment of CRISPR RNA delivered by LipofectamineMessangerMax or βN2-40. (**b)** Quantification of viral infection and (**c)** viability after therapeutic treatment of CRISPR RNA delivered by LipofectamineMessangerMax or βN2-40

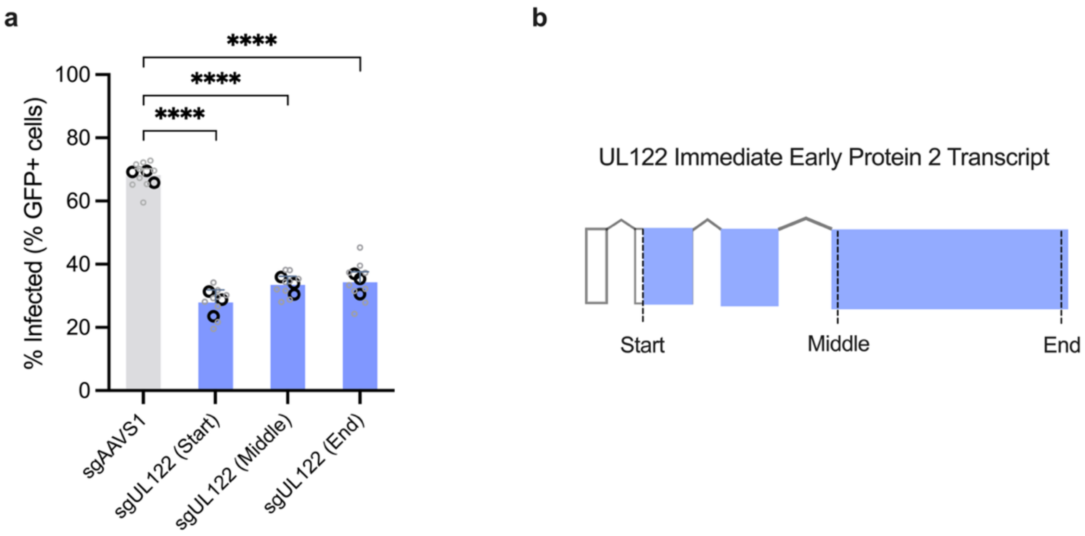


**Supplementary Fig. S13: Antiviral activity of CRISPR RNA LNP targeting different regions of the UL122 gene. (a)** % GFP+ cells after prophylactic and therapeutic CRISPR RNA LNP treatment targeting the start, middle, and end of the UL122 transcript. **(b)** Schematic depicting the relative location of the antiviral target. The diagram is not to scale. Black and gray data points denote biological and technical replicates, respectively. Statistical analysis was performed on biological replicates using one-way ANOVA for **a.** ****$P \leq 0.0001$

### 3. Additional References


1. Dunn, W. *et al.* Functional profiling of a human cytomegalovirus genome. *Proceedings of the National Academy of Sciences* **100**, 14223–14228 (2003).
2. Chambers, J. *et al.* DNA Microarrays of the Complex Human Cytomegalovirus Genome: Profiling Kinetic Class with Drug Sensitivity of Viral Gene Expression. *J Virol* **73**, 5757–5766 (1999).
3. Rozman, B. *et al.* Temporal dynamics of HCMV gene expression in lytic and latent infections. *Cell Rep* **39**, 110653 (2022).
4. Zhang, T. *et al.* Thioxothiazolo[3,4-a]quinazoline derivatives inhibit the human cytomegalovirus alkaline nuclease. *Antiviral Research* **217**, 105696 (2023).
5. Wang, D. & Shenk, T. Human Cytomegalovirus UL131 Open Reading Frame Is Required for Epithelial Cell Tropism. *J Virol* **79**, 10330–10338 (2005).
6. Nightingale, K. *et al.* High-Definition Analysis of Host Protein Stability during Human Cytomegalovirus Infection Reveals Antiviral Factors and Viral Evasion Mechanisms. *Cell Host & Microbe* **24**, 447-460.e11 (2018).
7. Marshall, E. E., Bierle, C. J., Brune, W. & Geballe, A. P. Essential Role for either TRS1 or IRS1 in Human Cytomegalovirus Replication. *J Virol* **83**, 4112–4120 (2009).
